# Initial niche condition determines the aging speed and regenerative activity of quiescent cells

**DOI:** 10.1101/2022.09.06.506737

**Authors:** Qi Liu, Nan Sheng, Zhiwen Zhang, Chenjun He, Yao Zhao, Haoyuan Sun, Jianguo Chen, Xiaojing Yang, Chao Tang

## Abstract

Quiescent cell ages with decline in both the survivability and regenerative activity. While most cellular quiescence/ageing research have focused on the survivability and from the population level, the question how the regenerative activity change with the quiescence time (i.e., chronological age) has rarely been addressed quantitatively. In this work, we systematically measured both features in ageing quiescent fission yeast cells at single cell level. We found that the regenerative activity declines linearly before survivability decline and the cellular chronological ageing speed is predetermined by the initial niche condition. Moreover, this linear ageing behavior is robust under various niche conditions and follows a common ageing trajectory in terms of gene expression. Furthermore, initial calorie restriction was found to improve not only the survivability but also the later regenerative activity. Our results reveal a continuous diverse spectrum of quiescence depth and ageing plasticity.

## INTRODUCTION

Cellular quiescence is a reversible non-proliferating cell state, in which cell ceased growth and division but retained regenerative capability to resume proliferation^1–3^. Endowed with this ability, quiescent cells function widely in many biological processes both physiologically and pathologically, e.g., tissue regeneration^4^, seeds germination^5^, plants longevity^6^, chronic infection^7^, and cancer recurrence^8^. Functionally, this regenerative capability can be dissected into two aspects, survivability and regenerative activity, which depicts how long a quiescent cell can survive and how fast it can recover from quiescence, respectively. During quiescence, the cell undergoes chronological ageing simultaneously. While the survivability has been extensively studied in terms of chronological lifespan in stationary yeast cells at the population level^9–14^, the regenerative activity has rarely been addressed quantitatively^15^. Considering many regenerative functions require not only long-term survival but also responsive recovery of quiescent cells, it is fundamentally important to understand at the single cell level how the regenerative activity of quiescent cell evolves during cellular chronological ageing and what factors influence this.

To avoid the heterogeneity and rapid death in the traditional stationary quiescence model, the stationary cells (ST cells hereafter) in budding yeast^16,17^, we used a long-survived and more homogeneous unicellular quiescence model, the nitrogen-starvation induced quiescent fission yeast *Schizosaccharomyces pombe* (Q_-N_ cells hereafter)^18,19^. By quantitatively study the evolution of single cell regenerative capability during chronological ageing under diverse niche conditions, we found robust linear ageing behaviors and a common ageing trajectory. Moreover, we found the cell ageing speed is determined by the initial niche condition and calorie restriction effect exists not only in the survivability but also in the regenerative activity.

## RESULTS

### Robust linear ageing behaviors at single cell level

To get a comprehensive view of how the regenerative capability of quiescent cells changes during cellular chronological ageing, we simultaneously measured the survival rate and recovery time (T_recovery_) of Q_-N_ cells under a wide range of nutrients availability with different initial suspension cell density spanning from optical density (OD) 0.15 to 1.0 (Figure 1A). The traditional ST condition was also included for comparison. The survival rate and T_recovery_ were used to quantify the survivability and regenerative activity, respectively, and the age of Q_-N_ cells was simply represented by the starvation duration (T_starvation_,). Note that the T_starvation_ includes a six-hour entering phase^18,19^, and short T_recovery_ indicates high regenerative activity.

**Figure 1.**
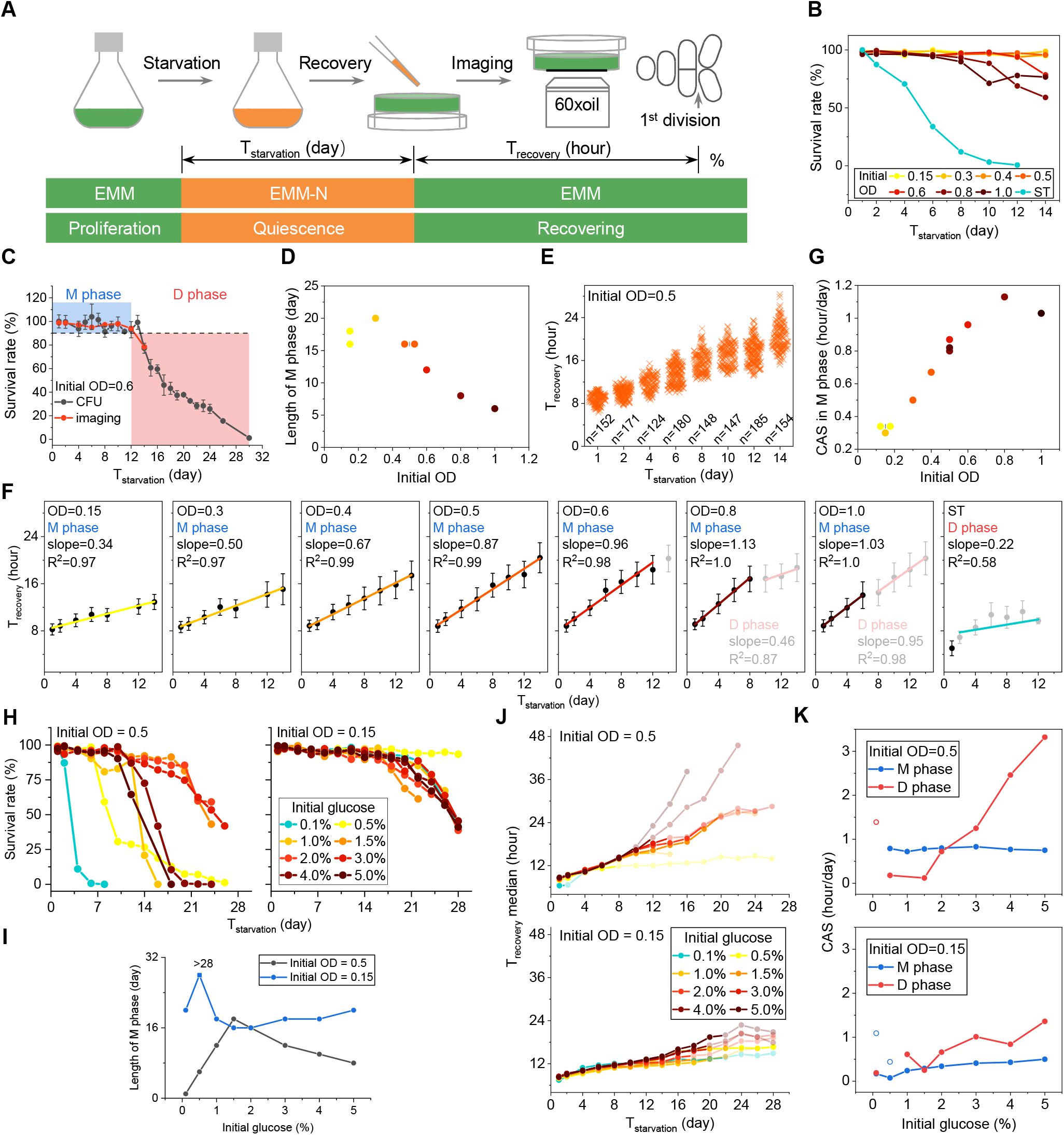
Quiescent cell regenerative capability and aging behaviors under various OD and glucose conditions. **(A)** Schematic of experimental design for measuring regenerative activity (T_recovery_) and survivability at single cell level by time-lapse imaging. **(B)** Survival rates under various initial ODs. **(C)** The schematic description of M phase (survival rate≥90%, blue) and D phase (red) from survival rate curve measured by sing-cell imaging (orange line), as exemplified by the condition of initial OD=0.6. Survival rate measured by CFU counting (black line, data are means ± SEM and represented four repeats) is included for comparison. **(D)** Correlation between the length of M phase and initial OD. **(E)** T_recovery_ vs. T_starvation_ with initial OD=0.5. Each cross represents a single cell and cell numbers are indicated. **(F)** Linear fitting of T_recovery_ vs. T_starvation_ data during M phase. Data are means ± SEM and cell numbers are listed in Table S1. Gray dots represent D phase. **(G)** The chronological ageing speed (CAS, defined as the linear fitting slope in **F**) under various initial ODs. **(H-K)**, Measured survival rate (**H**), length of M phase (**I**), T_recovery_ median (**J**), and CAS (**K**) under various glucose conditions under initial OD of 0.5 and 0.15. Cell numbers related with **B**, **F**, **H** and **J** are listed in Table S1 and S2.

Quantification from single cell imaging data showed that the survivability of quiescent cells was declining with age in an OD-dependent manner, cells with lower-OD survived longer (Figures 1B–1D, Table S1). After two-weeks’ starvation, Q_-N_ cells still maintained full survivability under ODs below 0.5 while dropped visibly under ODs above 0.6 (Figure 1B). Considering the quiescence state may be altered by the nitrogenous substances released from dead cells, we further divided the whole process into a maintaining phase (M phase, survival rate ≽ 90%) and a decaying phase (D phase) (Figure 1C). Note that, here, for simplification, the M phase as defined did not exclude the initial entering phase (E phase) during which the cells reduce their size through two rounds growth-absented division and shutdown the proliferation program^19,20^. The length of M phase was negatively correlated with the cell density (Figure 1D). The survival rate measured by imaging showed a very good agreement with the measurements by traditional CFU (colony formation unit) counting (Figure S1A). Generally, Q_-N_ cells survived much longer than ST cells (Figure 1B), consisting with previous studies^21,22^. This longevity of Q_-N_ cells enabled us to trace the ageing behavior from the single cell level in terms of regenerative activity during quiescence maintainment.

Single cell T_recovery_ measurements showed that the regenerative activity is progressively declining with age in all conditions including ST with an OD dependent manner (Figures 1E–1G). Generally, T_recovery_ for higher-OD cell was larger and Q_-N_ cells recovered slower than ST cells (Figures S1B and S1C). Intriguingly, T_recovery_ increased *linearly* with T_starvation_ during the M phase under all conditions except for the ST case in which the M phase was too short to tell (Figures 1E and 1F). Moreover, the linearity was nearly perfect with the R^2^ for linear regression was almost equal to one (Figure 1F). The linear fitting slope, which is the T_recovery_ changing rate, was defined as “chronological ageing speed” (CAS) for quiescent cells. We found that the CAS was positively correlated with cell density (Figure 1G). However, there seemed to be an upper limit for both survivability and CAS under the standard EMM-N condition (Figures 1D and 1G). Repeated experiments showed that the linear ageing behavior of quiescent cells was robust (Figures S1C–S1E). We concluded that quiescent cell ages linearly before survivability decline and its ageing speed is determined by the initial niche condition.

### Aging dynamics under various glucose conditions

Our previous results, that the higher recovery activity under lower OD and short survivability under glucose-exhausted ST condition, indicated that OD-dependent ageing behavior may be due to the different nutrients availability for individual cells. Considering that glucose might be the most important nutrient during nitrogen starvation and that has been reported to be a significant pro-ageing factor affecting the chronological lifespan of ST cells in both budding and fission yeast^12,21^, we next investigated the ageing behavior of Q_-N_ cells under various glucose concentrations (Figures 1H–1K). We applied a wide range of initial glucose concentrations spanning from 0.1 to 5.0% under two representative initial ODs, OD 0.5 and 0.15 (Figure S2A).

The survivability was significantly influenced by the glucose availability in an OD dependent manner (Figures 1H and 1I, Table S2). Q_-N_ cells under higher glucose concentrations were generally live longer, especially under high OD conditions (Figure 1H), suggesting long-term survival at quiescence state requires energy supply. However, the influence of glucose on survivability was not monotonic, as shown by the curve of M phase length in Figure 1I (Figures S2B and S2C). There existed an optimal glucose concentration for the survival of Q_-N_ cells, above which excess glucose was detrimental. The optimal glucose concentration for OD=0.15 was around 0.5%, which was much lower than the 1.5% for OD=0.5 (Figure 1I), suggesting again excess glucose is detrimental. We note that the maintained survivability under conditions above the optimal glucose concentration under OD 0.15, as indicated by the slight lifting in the M-phase length curve in Figure 1I, was due to the slight expansion of quiescent population which will be discussed later. The non-monotonic glucose effect on survivability demonstrated that glucose also promotes ageing in long-survived Q_-N_ cells.

For the regenerative activity, consistent with previous experiments with different initial ODs (Figure 1F), T_recovery_ increased linearly with T_starvation_ before survivability decline in all cases with various initial glucose concentrations (Figures 1J, S3A and S3B). However, unexpectedly, while the T_recovery_ increased significantly with the increase of initial OD, there was barely a change with the increase of initial glucose concentration under fixed OD (Figures S3C and S3D). Indeed, the T_recovery_ under all glucose concentrations had not diverged until at the end of M phase (Figure 1J, non-transparent dots). Consistent with this, the CAS was almost the same in the M phase, especially for the OD=0.5 case (Figure 1K, blue lines). Thus, the effect of initial OD on regenerative activity during M phase cannot be explained solely by glucose. Interestingly, after entered the D phase, T_recovery_ increased significantly with the increase of glucose concentration under both ODs (Figure 1J, transparent dots), suggesting glucose actually exerted a profound influence on the regenerative activity. In addition, it is worth noting that, normally the CAS should decrease a bit in the D phase, since many substances were released into the environment after cell death, as we observed before (Figure S1A, OD=0.8 and 1.0). However, here as the system entered into the D phase, the CAS increased significantly with glucose concentration above the optimal concentration for both ODs (Figure 1K, red lines), suggesting glucose promotes ageing on regenerative activity as well. Taken together, these results suggested pro-ageing effect by glucose exists not only on the survivability but also on the regenerative activity in quiescent cells.

### New anti-ageing effect by CR

Calorie restriction (CR) has been widely reported to extend chronological lifespan in various species^12,21,23,24^ However, little is known about the CR effect on the regenerative activity of quiescent cells. We have included two CR conditions (0.1% and 0.5% initial glucose) for the two initial ODs in our previous measurements (Figure S2A). Consistent with the well-known effect of CR, we also observed the pro-longevity effect in Q_-N_ cells under the CR conditions in both ODs (Figure 1H, yellow lines). Note that the rapid loss of viability under OD 0.5 with 0.1% glucose was due to the glucose deficiency for individual cells under high cell density.

For the regenerative activity, we found that in the young Q_-N_ cells in M phase, there was no significant difference between CR and non-CR conditions (Figure 1J, non-transparent dots). However, after entered D phase, aged Q_-N_ cells under CR conditions recovered significantly faster than under non-CR conditions (Figure 1J, transparent dots), suggesting CR has anti-ageing effect on the regenerative activity of quiescent cells. This trend is systematically consistent with the previously found pro-ageing effect of high-concentration glucose on the regenerative activity in the D phase. Although regenerative activity can also be improved by substrates released from dead cells during D phase, it either sustained short (Figure S1A, OD = 0.8 and 1.0) or appeared very late (Figure S3B, red dotted lines). The continuous high regenerative activity from the beginning of D phase under the CR condition of 0.5% glucose with OD=0.5 was very likely own to the CR effect. It is noteworthy that there were two different CAS during M phase under the CR conditions of OD=0.15, and the first one was larger than the second (Figure 1K, blue circles; Figure S3B, blue lines). While the second smaller CAS during M phase can own to the CR effect, the beginning larger CAS was probably due to a stress response to the very low glucose concentration. Consistent with this, the CAS under the CR condition with OD=0.5 was also high (Figure 1K, red circle).

Taken together, CR improves not only the survivability but also the regenerative activity of quiescent cells. However, the CR effect on regenerative activity appeared slowly not obvious until the population stars to decay.

### Ageing dynamics under other nutrients conditions

Since the effect of initial OD on regenerative activity cannot be explained solely by glucose (Figures S3C and S3D), we then asked whether other nutrients take part in it. By simultaneously adjusting the concentrations of vitamin and mineral elements (VM hereafter) in the EMM-N medium (see Methods), we investigated the influence of non-glucose nutrients on the regenerative capability of Q_-N_ cells under two representative ODs (Figure S4). Interestingly, increasing VM did not bring any significant effects but decreasing VM significantly impaired it (Figures S4A–S4C, Table S3). These results suggested that long-term survival in quiescent state requires not only glucose but also other nutrients. For the regenerative activity, increasing VM did not bring any positive or negative effect to the T_recovery_, while decreasing VM influenced T_recovery_ significantly, especially for OD=0.5 case (Figures S4D–S4F). Again, T_recovery_ was linearly increasing with T_starvation_ for all VM cases studied (Figures S4D and S4E). The CAS in M phase was maintaining unchanged under increased VM conditions but slowed down under decreased VM conditions (Figure S4F, blue lines). These results suggested that deficiency in VM nutrients may contributed to the change in regenerative capability and ageing speed. However, comparing with the significant effect of OD on T_recovery_ in M phase, the effect of VM was rather limited. Thus, the effect of initial OD on T_recovery_ cannot be simply explained by either glucose or VM. Considering cell density does not only influence the nutrient availability but may also bring other unknown complex effects to the global environment, the OD dependence of T_recovery_ and CSA could be a comprehensive effect of the whole niche condition.

### Correlation between regenerative activity and cell size

Although cell proliferation is ceased at quiescence state, we still observed cell size change under some conditions. The cell size increased significantly under OD 0.15 but almost unchanged under OD 0.5, and slightly decreased under OD 1.0 (Figures 2A and 2B), suggesting quiescent cells are metabolically active. It should be mentioned that, for the same OD, Q_-N_ cells under nutrients-sufficient conditions displayed no significant difference on cell size (Figure S5A). Considering cell size is a general reflection of the functional cellular components reserve which may influence regenerative activity, we then examined the correlation between T_recovery_ and quiescent cell size. Indeed, we found the T_recovery_ was negatively correlated with quiescent cell size under all conditions at all ages (Figure 2C). Within the same population, larger quiescent cells recovered faster than smaller ones. To quantitatively investigate the influence of the cell size on T_recovery_, we fitted T_recovery_ *vs*. size data with linear regression and defined the absolute value of the fitting slope as “size sensitivity” (Figure 2D), which quantifies how T_recovery_ depends on quiescent cell size. For most ODs, the size sensitivity was progressively increasing with chronological age and T_recovery_ was more depended on quiescent size at higher OD (Figures 2E and S5B), suggesting the intracellular components reserve in quiescent cells is getting more important at older age especially for higher OD conditions. Interestingly, Q_-N_ cells under OD 0.15 seemed to maintain a constant size sensitivity (Figures 2E and S5B), suggesting that they relied less on their own intracellular reserve, probably due to the relatively abundant niche resource under low cell density.

**Figure 2.**
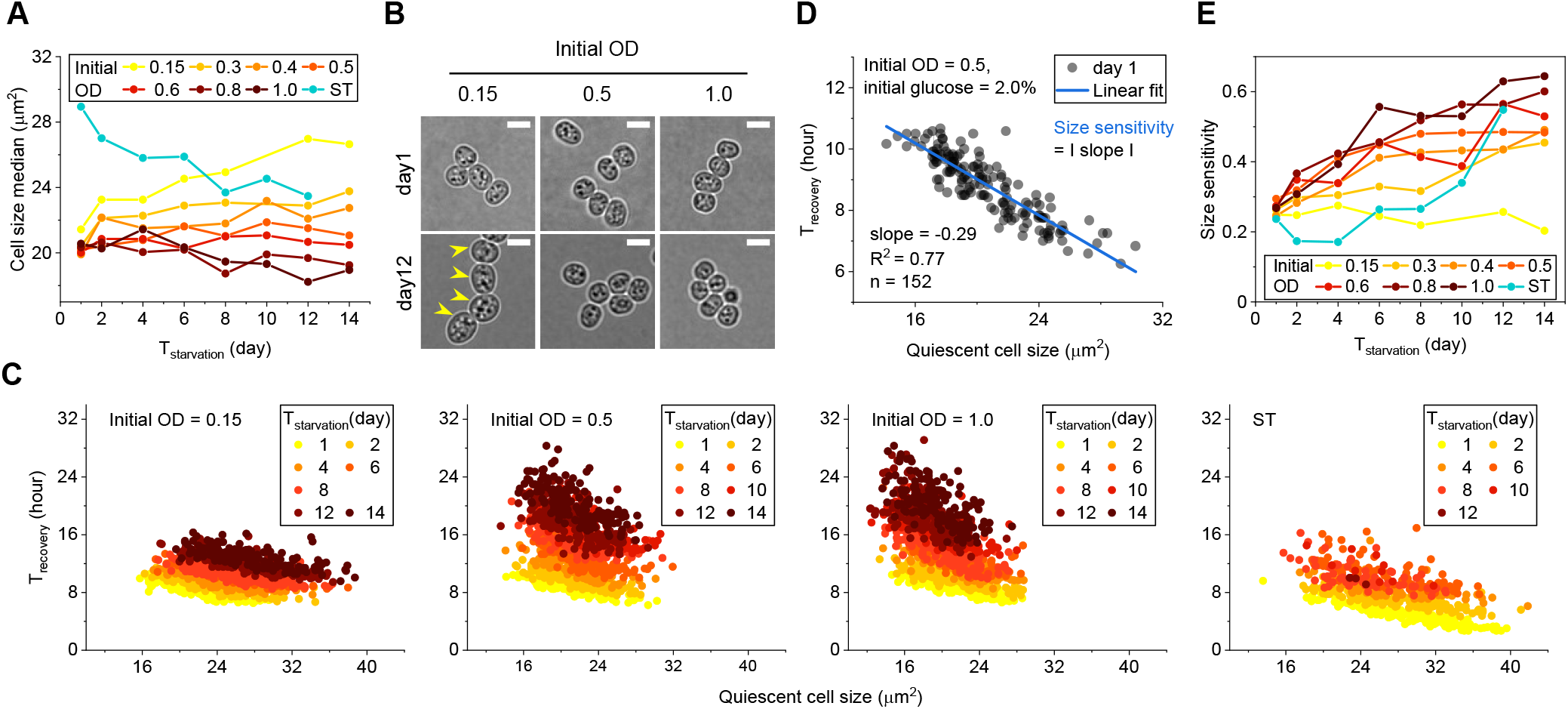
Correlation between T_recovery_ and quiescent cell size. **(A)** Comparison of cell size median of quiescent cells under various initial ODs. (**B**) Microscope images of typical Q_-N_ cells at day 1(up) and day 12 (bottom) under three initial ODs. Yellow arrows indicate size-increased cells. The scale bar is 5μm. (**C**) T_recovery_ was negatively correlated with quiescent cell size under all conditions, exemplified by the initial ODs of 0.15, 0.5, 1.0 and ST condition. Each dot represents a single cell. (**D**) Linear fitting the negative correlation between T_recovery_ and quiescent cell size, exemplified by the initial OD of 0.5. The absolute value of the fitting slope is defined as “size sensitivity”. Data condition and fitting results are indicated. **(E)** Comparison of size sensitivity of ST and Q_-N_ cells under various initial ODs. Each dot in **C** and **D** represents a single cell. Cell numbers for data in **A** and **C** are listed in Table S1.

### Intracellular change during ageing

To understand physiologically how intracellular functional components may influence the regenerative activity of quiescent cells during ageing, we then investigated the change of several intracellular components, which include mitochondrion, nuclear, vacuolar, ribosome, cytosol metabolic enzyme, lipid droplet, and cell cycle regulators. We labelled some key proteins with fluorescent tag and imaged their morphology and location at different age under two representative initial ODs (Figures 3A and S6A).

**Figure 3.**
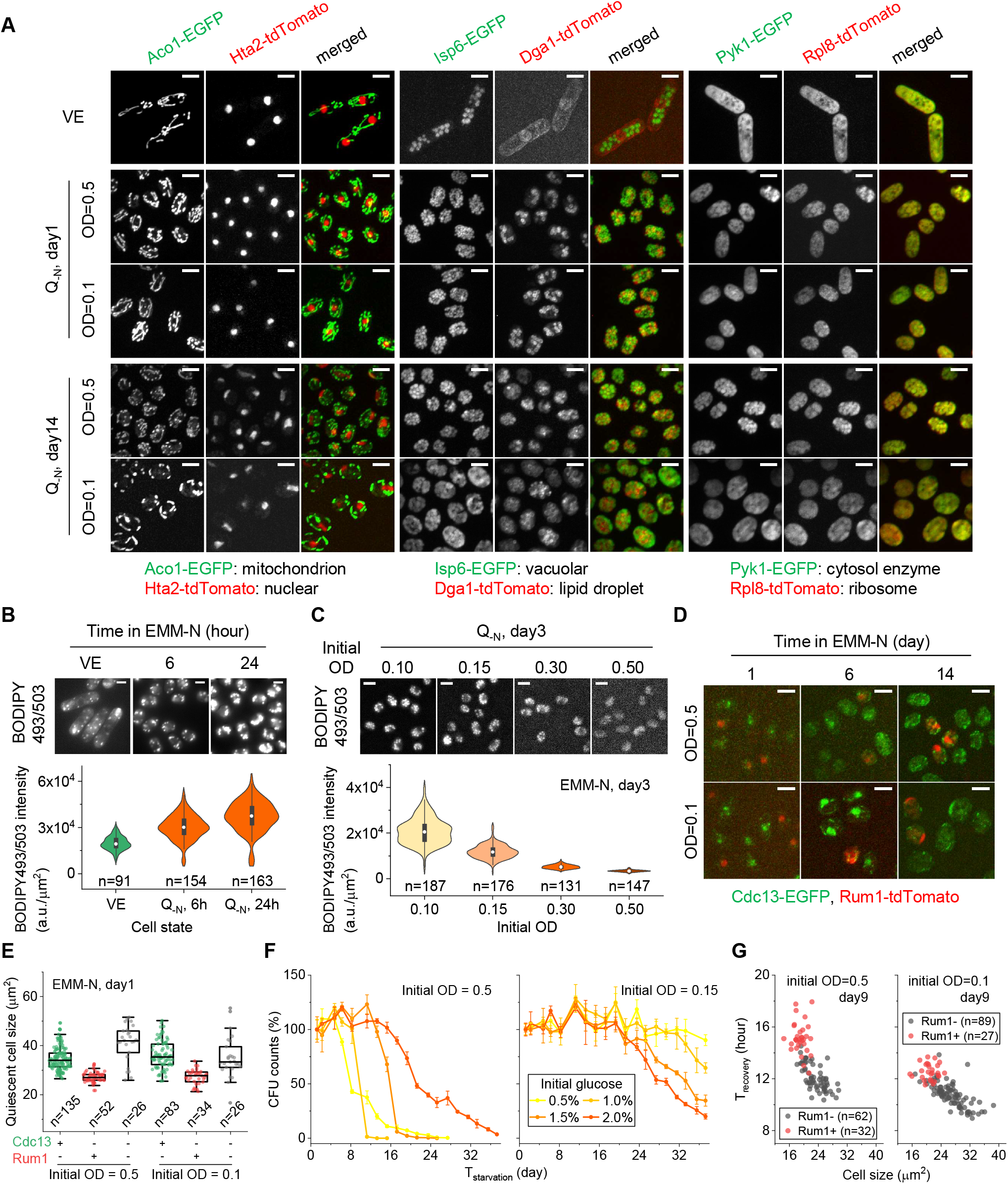
The change of intracellular functions during quiescence development and ageing. **(A)** The fluorescence microscope images of presentative functional proteins at different ageing stages under initial OD of 0.5 and 0.1. Proteins are tagged with fluorescence protein and their representative functions are indicated. **(B-C)** Comparison of lipids accumulation at different quiescence time under the initial OD 0.5 (**B**) and under various initial ODs at day 3 (**C**). Up and bottom panel in **B** and **C** are microscope image of lipids stained with BODIPY493/503 and lipids quantification respectively. Note that cells in **C** were washed by 1xPBS after staining. **(D)** Fluorescence microscope images of cell cycle regulators at three quiescence ages under initial OD of 0.5 and 0.1. **(E)** Comparison of quiescent cell size for three subpopulation of Q_-N_ cells with different cell cycle signals at day 1 under two initial ODs. “+” and “–” represents positive or negative in the appearance of nuclear located signals respectively. **(F)** Survival rates measured by CFU counting of Q_-N_ cells under four glucose conditions with the initial OD of 0.5 (left) and 0.15 (right). **(G)** Comparison of T_recovery_ of Q_-N_ cells with (Rum1+) or without (Rum1-) Rum1-tdTomato signal. Each dot in **E** and **G** represents a single cell. The scale bar is 5μm.

Comparing with vegetatively proliferating cells (VE cells hereafter), Q_-N_ cells displayed obvious mitochondrion fission (Aco1-EGFP), nucleus condensation (Hta2-tdtomato), vacuolar expansion (Isp6-EGFP), cytosol and ribosome granulation (Pyk1-EGFP and Rpl8-tdTomato), and lipid droplets accumulation (Dga1-tdTomato) under both ODs (Figure 3A), suggesting a global organelle and metabolic remodeling during the quiescence development. Again, we observed quiescent cells with obvious size increasement under lower OD (Figures 3A and S6A). Mitochondrion fission and nucleus condensation are generally correlated with low energy state and inactive transcription. We found the extent of mitochondrion fission at old age was stronger in OD=0.5 than that in OD=0.1 (day14, Figure 3A), suggesting relatively lower energy level under higher OD.

Vacuole is an acidic organelle with degradation and storage functions^25^. The vacuole protease Isp6p in fission yeast was reported to be specifically induced by nitrogen starvation and participated in autophagy which functions in nitrogen recycling by bulk degradation of intracellular proteins^26–29^. We found the number of vacuoles was significantly increased at the very early age (day1, Figure 3A), their size was getting bigger as cell was getting older, and the size of vacuoles was even bigger under lower OD (day14, Figure 3A), suggesting higher bulk degradation and nutrients storage capability under lower OD. It is worth mentioning that, in both ODs, the cytosol glycolytic enzyme Pyk1-EGFP and ribosome protein Rpl8-tdTomato, which are both spreading in the VE cytosol, also displayed vacuole-like structures (Figure 3A). Indeed, both Pyk1-EGFP and Rpl8-tdTomato were well co-localized with Isp6-containing vacuoles (Figure S6A), suggesting the delivering of cytosol enzymes and ribosome into the bulk degradation and nitrogen recycling system.

Lipid droplets (LDs) are the storage organelle of neutral lipids such as triacylglycerols (TAG) and play central function in lipid and energy homeostasis^30^. Dga1 is a triacylglycerols synthase in fission yeast and its deletion reported to cause TAG deficiency and apoptosis in ST cells^31^. While in VE cells Dga1-tdTomato predominantly located in the nuclear and peripheral membrane, in Q_-N_ cells it was significantly up-regulated and largely located in LD membrane (Figure 3A), suggesting increased triacylglycerols synthesis. LD-staining by BODIPY493/503^32^ confirmed the accumulation of lipids as cell entered into quiescence (Figure 3B, Movie S1). Furthermore, we found quiescent cells accumulated more lipids under lower ODs (Figure 3C). It should be mentioned that fission yeast cells form basal level of LDs at proliferating state (Movie S2). However, these basal level lipids in LDs were quickly consumed in ST cells (Figure S6B).

Fission yeast cell cycle was driven by a major cyclin Cdc13^33–35^ and blocked before G1 under poor nutrition by a negative regulator Rum1^36,37^ It was reported that fission yeast experience two cell divisions to arrest cell growth and exit cell cycle before quiescence commitment^19^. Interestingly, after quiescence commitment, the nuclear-located Cdc13-EGFP signal can still be seen in most cells, and only a small subgroup of Q_-N_ cells displayed Rum1-tdTomato signal (day1, Figure 3D and 3E). In addition, Cdc13-EGFP signal mainly existed in bigger Q_-N_ cells while Rum1-tdTomato signal was exclusively expressed in smaller ones (Figure 3E). After six days’ nitrogen starvation, the nuclear-located Cdc13-EGFP signal was weakened under OD 0.5 but still sustained under OD 0.1 (Figure 3D), suggesting Q_-N_ cells under lower cell density retained longer cell division activity. Consistent with this, CFU counting experiments showed that the quiescent population was obviously expanded under lower cell density condition (Figure 3F), suggesting some quiescent cells displayed a further cell division during later quiescence state. However, further into the nitrogen starvation, the Cdc13-EGFP signal was either disappeared or compacted into the condensed nucleus under both ODs (day14, Figure 3D), suggesting the cell cycle program was eventually inactivated at old age. Generally, Q_-N_ cells with positive Rum1-tdTomato signal recovered slower than Rum1-tdTomato negative cells (Figure 3G). These data demonstrated that quiescent cells are not necessarily existed the cell cycle after quiescence commitment, consistent with the previous study that quiescence development is not driven by cell cycle signals^38^. Depending on the OD and cell size, quiescent cells may shut down the cell cycle program slowly during ageing.

Taken together, these results suggested that the overall cellular organization and metabolism were remodeled during the quiescence development. However, the remodeling extent was greatly dependent on the niche condition.

### Common ageing trajectory

In order to explore the whole ageing process of quiescent cells at the molecular level, we further performed RNA-seq on Q_-N_ cells under four representative conditions (group HS OD=0.5, 2% glucose; group HH OD=0.5, 5% glucose; group LS: OD=0.15, 2% glucose; group LR: OD=0.15, 0.1% glucose) at different time points (3hr, 6hr, 9hr, 12hr, 1d, 4d, 7d(8d), (10d), 14d, (15d), 21d). Note that group HS was taken as the standard condition for Q_-N_ cells and group LR for CR condition.

We put all the samples together and calculated the correlation coefficients of overall gene expression between all pairs. They clustered into six clusters (Figure 4A). Interestingly, the six clusters turned out to be highly correlated with chronological age. The first and second clusters included all first day samples for all cases. The third and fourth clusters corresponded to maintaining phase samples, and the fifth and sixth clusters corresponded to the decaying phase samples. The only exception is that the late maintaining phase samples under the CR condition were clustered into the fifth cluster (Figure 4A, group LR 7d, 10d and 14d). These results indicated that the ageing processes under non-CR conditions may follow a similar trajectory. We further performed the principle component analysis (PCA) with all samples, and the result confirmed that non-CR conditions followed the same trajectory but with different evolving speed (Figures 4B and 4C). Consistent with the previous finding that CAS increases with initial OD (Figure 1G), higher OD indeed had a higher evolving speed along the “ageing trajectory” in terms of the global gene expression (Figure 4C: PC2 group HS (rose red) vs. group LS (blue)), while no significant difference on evolving speed was found with different glucose concentrations (Figure 4C: PC2 group HS (rose red) vs. group HH (green)).

**Figure 4.**
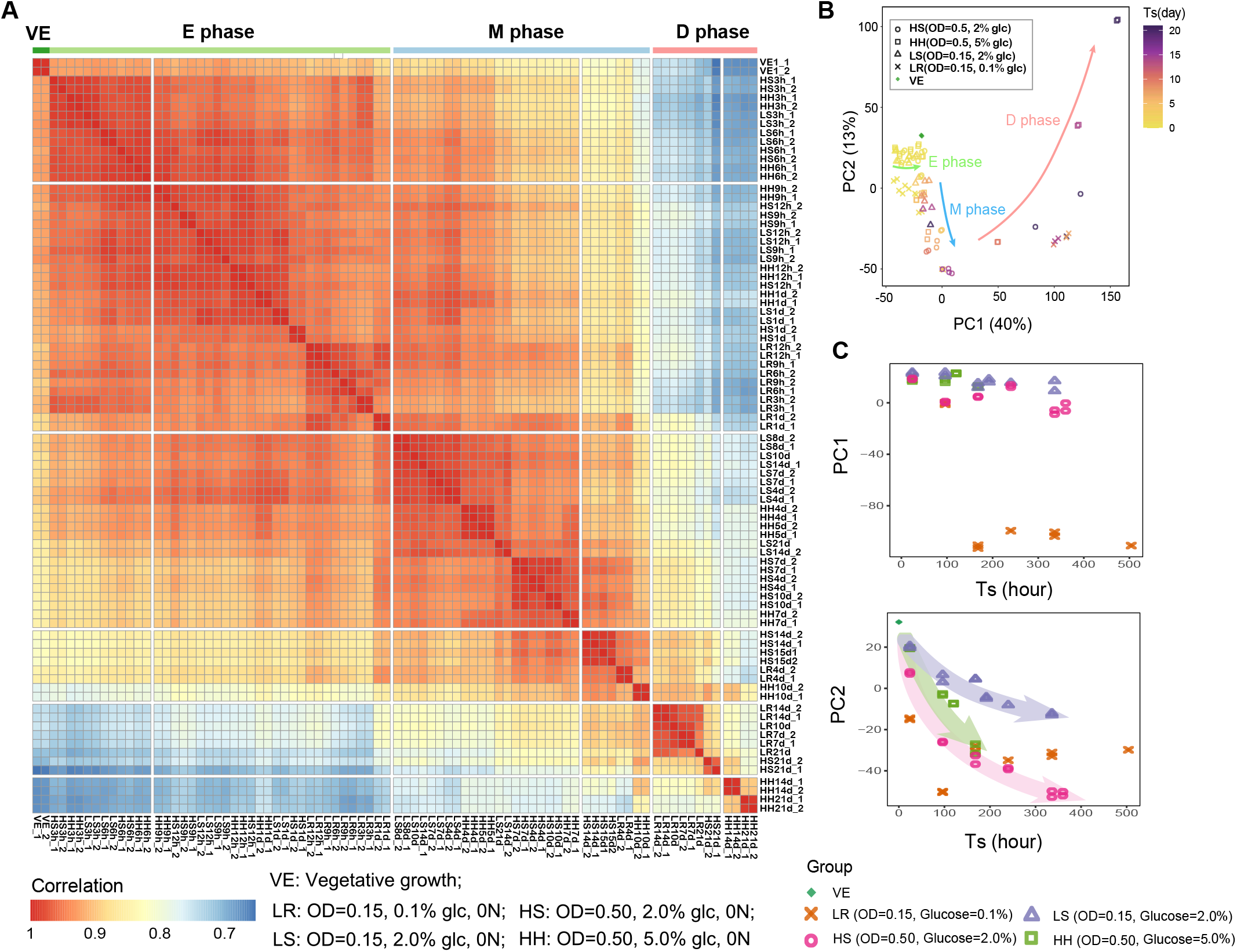
Quiescent cells under various niche conditions follow a common ageing trajectory. **(A)** Heatmap of correlation coefficients of overall gene expression between all samples. Color bar represents Pearson’s correlation coefficient. Group conditions are indicated. **(B)** Principle Component Analysis (PCA) of the global gene expression pattern of all samples. Each dot represents a single sample. Color bar indicates starvation time (T_s_). Shapes indicate different groups. Green, blue, and red guide lines indicate the general ageing trajectories during E, M, and D phase, respectively. **(C)** PC1 and PC2 of samples during M phase along the T_s_.

### Dynamic gene expression along the ageing trajectory

Since all results indicated the existence of a common characteristic ageing trajectory, we next investigated the gene expression changing pattern along this trajectory. The sequential change along the trajectory was reflected in 11 gene clusters that exhibited different temporal dynamic patterns (Figure 5A). It is worth noting that after clustering, gene expression was found to be divided into four distinct stages along the quiescence development: VE state, entering phase (E phase), M phase and D phase (Figures 4A and 5A), suggesting again the characteristic gene expression in different cellular states and ageing process.

**Figure 5.**
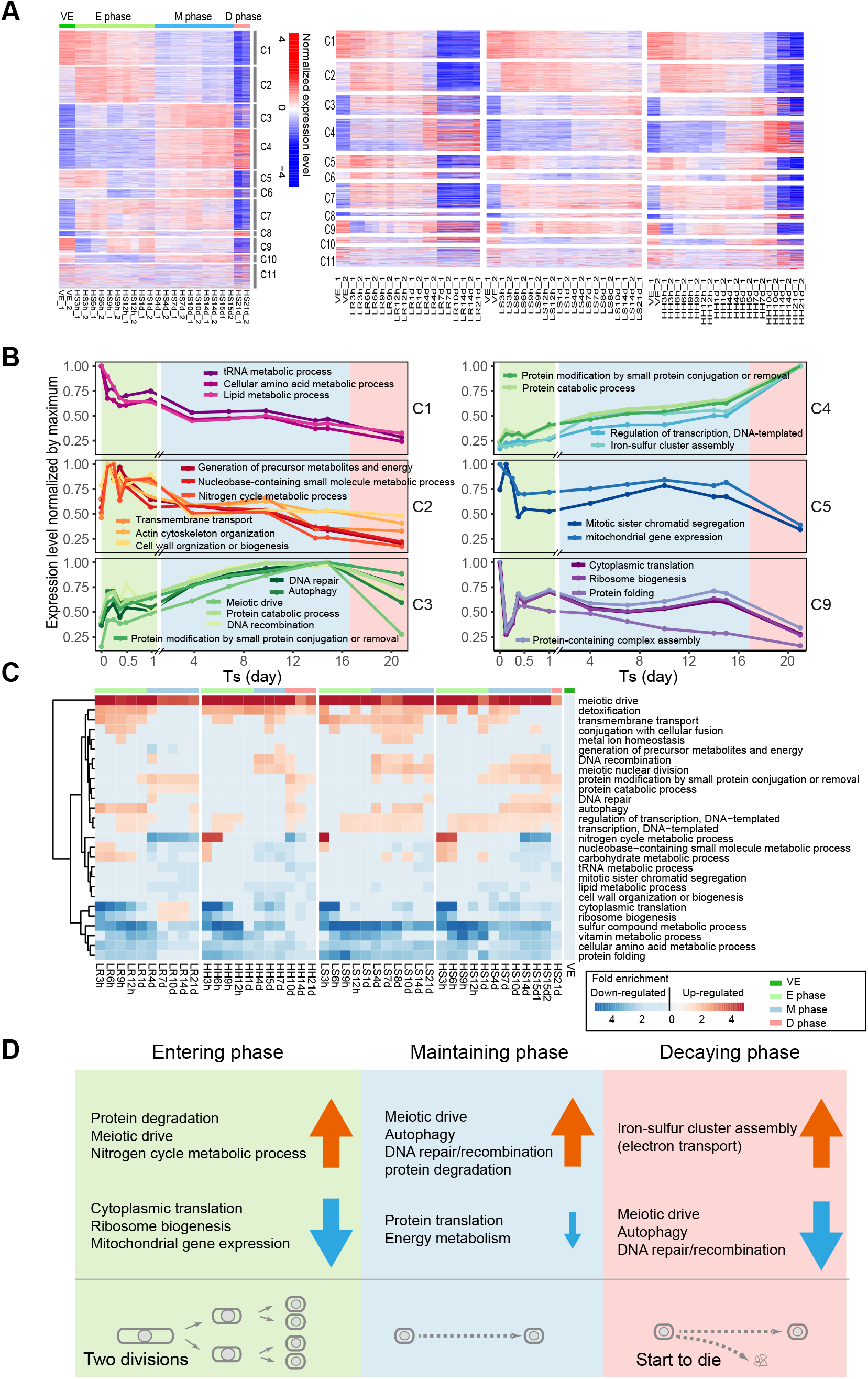
Dynamic gene expression along the ageing trajectory. **(A)** Gene expression profile during quiescence development and cellular ageing under four group conditions. Color indicates the normalized gene expression level. Up- and down-regulations were shown in red and blue respectively. Sample names with time were shown on the bottom, developmental and ageing stage were shown on the top. Genes were clustered to 11 clusters by hierarchical clustering as shown on the right. **(B)** Expression dynamics of enriched gene functional modules within clusters identified in **A** under the HS group condition. **(C)** Heatmap representing significantly enriched GO slim terms (rows) under four group conditions (columns) at different quiescence state compared with proliferating state (VE). Color bar indicates the fold enrichment of GO categories. **(D)** Summarized quiescence developmental and ageing program under nitrogen starvation.

We further did the functional enrichment within different 11 clusters, and found different expression dynamics for different function modules during the ageing process (Figures 5B and 5C). Take the group HS for example (Figure 5B), after 24 hours’ nitrogen depletion, while the protein degradation and nitrogen cycle metabolic process were upregulated, the protein translation and energy metabolism were downregulated dramatically. Meanwhile, meiosis related genes were also upregulated. As cells entered into the M phase, besides the further reduction of the protein translation and energy metabolism, meiosis related pathway kept being upregulated, and as well as the autophagy and DNA repair and recombination related genes (Figure 5B). Then, in the D phase, the meiosis, DNA repair, DNA recombination and autophagy related genes were all downregulated, and the electron transport related gene were upregulated. The characteristic quiescence development and ageing program for Q_-N_ cells was summarized in Figure 5D.

We also looked into the difference between different glucose and OD conditions (Figure 5C). Under non-CR conditions, though the evolving speed of group HS and HH were similar during the M phase, we still found some difference between them. Compared to 2% glucose, the detoxification related genes were significantly upregulated during the whole E-M-D process in 5% glucose. On the other hand, the upregulation of autophagy related gene can be detected during the whole M and D phases in 2% glucose, while it can only be detected at two time points in D phase in 5% glucose. These results suggested pro-ageing effect again for higher glucose concentration. As for the OD conditions, compared to OD=0.5, we observed a significant higher level of transmembrane transport, cytoplasmic translation and ribosome biogenesis in OD 0.15 (Figure 5C), suggesting a higher energy state level and higher metabolic activity for the lower OD case.

It is worth noting that the ageing trajectory for CR condition (group LR) was significantly different from the other three non-CR cases (Figures 4B and 4C). Firstly, no obvious upregulation of nitrogen cycle metabolic process related genes (Figure 5C). Secondly, autophagy related genes were upregulated at the very beginning of the starvation, while for non-CR conditions the upregulation of autophagy was much later (Figure 5C). Thirdly and most importantly, the cytoplasmic translation and ribosome biogenesis were significantly upregulated in the M phase, which were even higher than that of VE state (Figure 5C). These differences jointly programmed CR cells to survive longer and recover faster than non-CR cells at older age.

### Quiescence depth and spectrum

Our results showed that both Q_-N_ and ST cells exhibited significant difference in T_recovery_ at different age under various niche conditions (Figures 1F and 1J), suggesting ageing quiescent cells can enter a large variety of states with different regenerative activity. These data strongly supported the viewpoint that cellular quiescence is not a static “sleeping beauty”^39^ state but evolving with time and having different depth^40,41^.

To quantify the depth of quiescence state, we introduced a dimensionless parameter named “quiescence depth” (QD) which was defined as T_recovery_/T_cycle_. T_cycle_ was the cell cycle time for VE cells and measured to be 2.5h on agarose dish (Figure 6A). For comparison, we also included VE cells for QD analysis, with the single cell division time of full cell cycles taken as their T_recovery_. While the QD distribution for VE cells showed a narrow range with median value 1.0, the QD distributions for both Q_-N_ and ST cells were much wider, indicating a wide spectrum of quiescence states (Figure 6B). Interestingly, the QD distribution for ST cells showed a sizable overlap with VE cells, while there was no obvious overlap between Q_-N_ and VE cells (Figure 6B). It is worth noting that, under different glucose conditions, Q_-N_ cells displayed remarkably different QD (Figure 6C). For example, for the OD=0.5 cases, to fully resume proliferation after 16 days’ nitrogen starvation, on average 5, 8, and 12 cycles’ long time were needed for Q_-N_ cells under 0.5%, 2.0% and 4.0% glucose, respectively (Figure 6C, Movies S3-S5).

**Figure 6.**
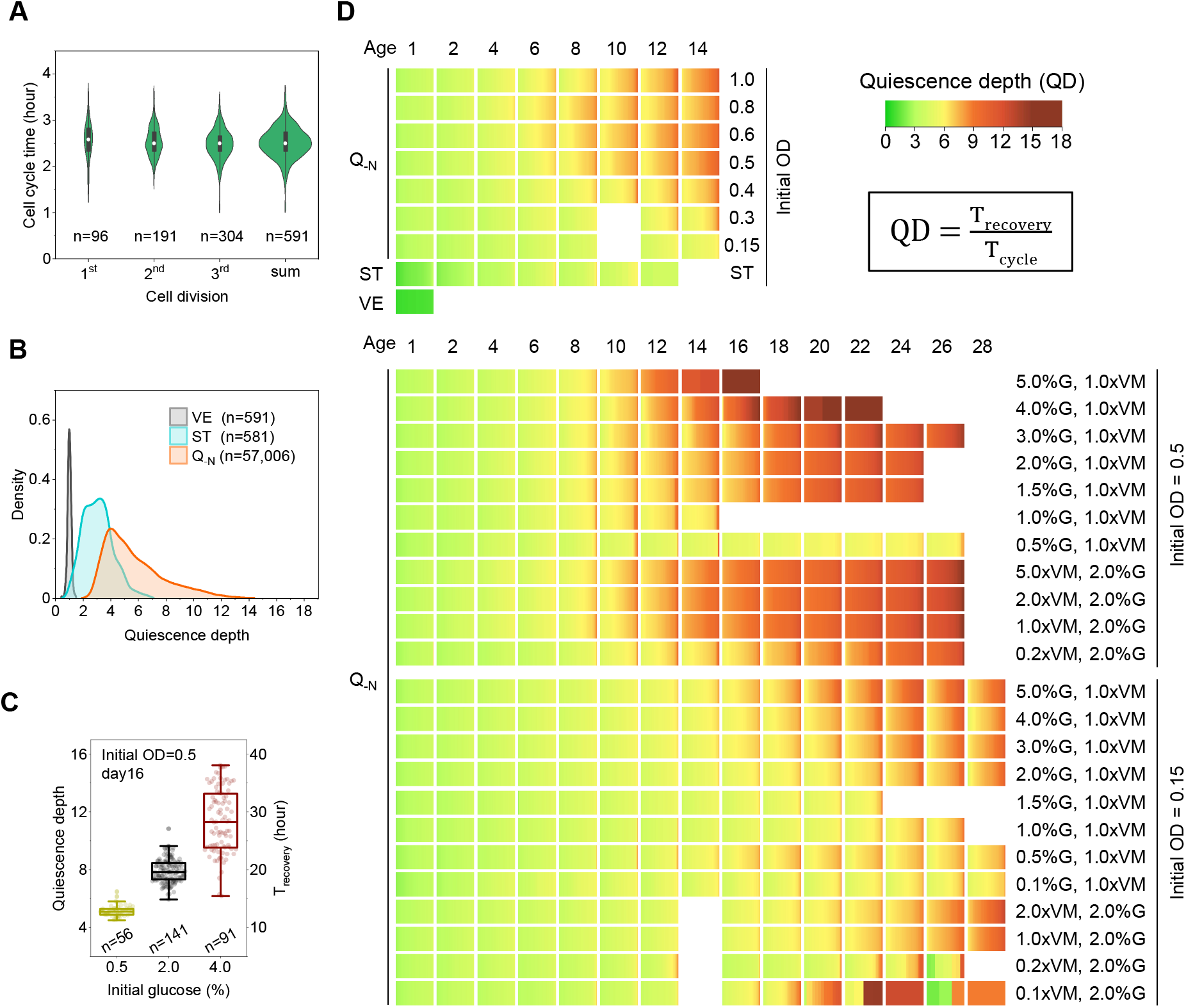
Ageing quiescent cells can enter a large spectrum of states with various quiescence depth and regenerative capability. **(A)** Distribution of cell cycle time of proliferating fission yeast cells measured on agarose dish. Three sequential divisions with full cell cycle were analyzed. Cell numbers were indicated. **(B)** The quiescence depth (QD) distributions of VE, ST, and Q_-N_ cells. Data for VE cells is the “sum” in **A**. Data for ST cells is the same with ST condition in Figure 1. Data for Q_-N_ cells is a combined data of all Q_-N_ conditions in Figure 1 and Figure S4. Total cell number for each category is indicated. **(C)** Comparison of QD of Q_-N_ cells at day 16 under three representative glucose conditions under the initial OD of 0.5. Data represented by boxplots overlapped with single cell data. Each dot represents a single cell with cell number indicated. **(D)** Visualized all single cell QD under total 32 niche conditions. Each block is an array of color strips representing single cell QD. The time point (age) and condition are indicated on the top and right of each panel. Cell numbers are listed in Tables S1 to S3.

The biological meaning for QD is that, the deeper the cell in quiescence state, the lower in regenerative activity and slower in resuming proliferation. To get an overview of the depth landscape for quiescent cells, we visualized all the single cell QD over various conditions (Figure 6D). In total, we analyzed 57,587 single quiescent cells with different chronological age and under 31 different niche conditions. Indeed, all the QD constituted a beautiful continuous spectrum (Figure 6D). The chronological age and niche condition made up the two extrinsic dimensions for the QD spectrum. On one hand, the QD was gradually increasing with age. On the other hand, the QD also relied on the niche condition such as cell density and nutrients availability. For example, the QD of quiescent cells could be modified by the niche changes in the decaying phase. Moreover, the spectrum property also existed in the same population within the QD block of quiescent cells with the same age. This QD spectrum together with survival curves illustrated that quiescence state is dynamic and plastic and ageing quiescent cells could exhibit dramatically different regenerative capability depending on the niche condition.

## DISCUSSION

The most essential function for quiescent cells is their regenerative capability. Quiescence cells also undergo chronological ageing, not only in the survivability but also in the regenerative activity. While most research concerning cellular quiescence and ageing mainly focused on the survivability and lifespan and from the population level, the efforts on the single cell regenerative activity and the ageing process *per se* were rare. In this work, we systematically and quantitatively measured both feature of ageing quiescent fission yeast cells from the single cell level. In conclusion, we found quiescent fission yeast cells age linearly in terms of regenerative activity before survivability decline under various niche conditions and follow a common ageing trajectory. Moreover, the cell ageing speed is determined by the initial niche condition and calorie restriction could improves not only the survivability but also the regenerative activity.

Cellular quiescence, also called cellular dormancy in many papers^2,3,7,8^, is commonly defined as a reversible non-proliferating cell state^1–3, 39^. Traditionally, this state is viewed as a unique state in comparison to the proliferation state. However, the essence of this state has been debated for a long time^39–41,42,43,44,45^. Our work from evolution perspective together with single cell quantification demonstrated that the nature of cellular quiescence is far from being unique and static and harboring diverse depth. Indeed, quiescence depth constitutes a wide range of continuous cell state spectrum (Figure 6D). Under different niche conditions, quiescent cells may exhibit dramatic difference in regenerative activity at the same age. Even though significantly different from the proliferation state in morphology and cellular activity, quiescent cells under some conditions and age may still harbor several gene expression features similar to it (Figure 3D and Figure 5). Our work provided an overall view on the whole quiescence landscape and may help to revisit the concept of cellular quiescence.

Quiescent cells also age chronologically while maintaining the regenerative capability. Chronological ageing and ageing rate have been mostly evaluated from the survivability aspect in terms of lifespan in both unicellular yeast models^9, 46, 47^ and multicellular organisms^23, 24, 48^ from either population level or species level. However, the lifespan of an organism is a comprehensive result of unsynchronized ageing processes in different cells within the same population or individual^49^. Thus, the lifespan itself is not the whole story of the ageing process. Understanding how a single cell ages requires direct measuring of the ageing-accompanied dynamic change in the characteristic features or functions of the cell, such as the regenerative activity of quiescent cells. Here in our work, from the perspective of regenerative activity which was quantified by the parameter T_recovery_, we directly described the ageing dynamics of quiescent cells under various niche conditions (Figures 1 and S4). From the T_recovery_, we unveiled the linear ageing behavior of quiescent cells and a new anti-ageing effect of CR which acts on the regenerative activity. All these new discoveries implied that under the current situation of lacking a common molecular marker for both cellular quiescence and cellular ageing, quantified parameters on the whole cell activity *per se* like T_recovery_ could be considered.

Remarkably, the T_recovery_ and cellular chronological ageing speed (CAS) under all glucose conditions were almost the same during most of the M phase (Figure 1K). The pro-ageing effect by glucose did not appear until very late at the end of M phase under the OD 0.15 and even only after entering the D phase under the OD 0.5 (Figure 1J). These results suggested that the pro-ageing effect by glucose is accumulative and its effect may require other nutrients. Interestingly, at late D phase, the increasing in T_recovery_ was eventually slowed down, probably due to the niche transformation by nutrients released from plenty of dead cells. This suggested that quiescence state is plastic and the regenerative activity of aged quiescent cells is changeable. Future tests would be needed to examine whether the aged regenerative capability of quiescent cells could be rejuvenated.

## MATERIALS AND METHODS

### Strains, plasmids, and media

All the fission yeast strains used in this study were congenic 972h- and listed in Supplementary Table 4. All fluorescence labeled genes were C-terminally tagged. Plasmids used for strain construction were listed in Supplementary Table S5. All constructs were confirmed by colony PCR. Oligonucleotides used in strain and plasmid construction were listed in Supplementary Table S6. Standard protocols were used throughout^50, 51^.

For construction of double-color strains LQF006, LQF068, LQF069, LQF072 and LQF079, plasmid PP004 and PP005 were used as template for the fluorescent cassette yEGFP-T_yADH1_-kanMX6 and tdTomato-T_yADH1_-natMX6 respectively. First genes were tagged with the fluorescent cassette yEGFP-T_yADH1_-kanMX6 by two-step PCR^51^ and the recombinant DNA were integrated into their endogenous locus. Successful transformants were selected by G418 (Sigma-Aldrich, Cat# A1720). Correctly integrated transformants were used for the second gene labeling. Second genes were tagged with the fluorescent cassette tdTomato-T_yADH1_-natMX6 by the same method and integrated into their endogenous locus. The double-color constructions were selected with G418 and nourseothricin (GOLDBIO, Cat# N500). For construction of the strain LQF001, the first gene Cdc13 was tagged with yEGFP-T_yADH1_-kanMX6 as previous. The second gene Rum1 was tagged with tdTomato and expressed under its native terminator T_Rum1_. The recombinant DNA Leftleg_Rum1_-tdTomato-T_RUM1_-natMX6-Rightleg_Rum1_ was first cloned into the plasmid PP008, then amplified by PCR from PP008 and integrated into the endogenous Rum1 site.

All plasmids were replicated in DH5α Escherichia coli. All constructs were confirmed by sequencing. The plasmid PP004 and PP005 were constructed by enzymatic digestion and ligation. Firstly, the kanMX6 cassette in the plasmid pDH3 was replaced with the natMX6 cassette from the plasmid pFA6a-natMX6 with BglII and EcoRI enzyme pair to generate the plasmid PP001. Then the CFP cassette in the plasmid pDH3 and PP001 were replaced by the yEGFP and tdTomato cassette from the plasmid PP002 and PP003 respectively with PacI and AscI enzyme pair. For construction of the plasmid PP008, the recombinant DNA Leftleg_Rum1_-tdTomato-T_yADH1_-natMX6-Rightleg_Rum1_ was prepared by two-step PCR and then cloned into the T vector pMD19 to generate the plasmid PP006. Then the native terminator of Rum1 (T_Rum1_) flanking with the AscI and BglII cloning site was prepared by PCR and then cloned into T vector pMD19 to generate the plasmid PP007. At last, the terminator T_yADH1_ in the plasmid PP006 was replaced by the T_Rum1_ terminator from the plasmid PP007 with AscI and BglII enzyme pair.

Synthetic media used for yeast culturing and quiescence maintaining are listed in Supplementary Table S7. All media were made according to the standard recipes^50^. YE agar plates with or without drug were used to select fluorescence constructions or grow CFU colonies respectively. EMM and EMM-N (EMM lacking NH4Cl) liquid media were used to prepare proliferating and quiescent cells respectively. EMM-N-C (EMM lacking NH4Cl and glucose) liquid medium was used to wash proliferating cells before re-suspending into EMM-N. EMM with 1.5% low melting temperature agarose (Lonza, Cat# 50080) was used for time-lapse imaging of quiescent cells’ recovery. EMM agarose containing 1ug/ml BODIPY493/503 (Invitrogen, Cat# D3922) was used for imaging the dynamics of lipids droplets in proliferating cells. All media ingredients related to EMM, EMM-N, and their variants were chosen as Sigma BioUltra grade.

### Microscope agarose dish preparation

Petri dish with a removable glass bottom (NEST Cat# 801002) was used to make EMM agarose dish. The original round glass bottom was removed and replaced by a square microscope cover glass (Fisher 12-542A) and sealed with Scotch tape (Cat# 810QC33). 20ml EMM liquid medium containing 1.5% (m/v) low melting temperature agarose was microwaved for no more than total 45 seconds and then cooled down to below 37°C under running water. For lipids droplets imaging, additional 20μl 1mg/ml BODIPY493/503 was add to the 20ml cooled EMM agarose. 2.7mL cooled liquid EMM agarose medium was immediately filled into the reconstructed petri dish and then thoroughly solidified at room temperature for about 3 hours. Before use, the agarose dish was inverted for 0.5 hour for easily tearing off the square cover glass without damaging the smooth agarose surface. The agarose dishes were freshly made before use on the experimental day. Just before loading cells, tear off the cover glass and dry the surface for several seconds at room temperature. Then drop 0.1μl cell suspension on the agarose surface and array all samples in the central region of the imaging area with a designed order. After each sample loaded, dry the sample spot for several seconds. Make sure all loaded spots are isolated and not cross diffused into each other. When all samples were settle down, record the time and then seal the agarose with a new clean cover glass and fix it with Scotch tape but leaving an imaging area as large as possible. It is essential to make sure that, (1) the agarose is thoroughly solidified, (2) the agarose surface is smooth after teared off the cover glass, (3) agarose surface is dried before loading each sample, otherwise the loaded cells will be cross diffused. After sophisticated, a ready-for-imaging microscope agarose dish loaded with up to 10 isolated samples can be made within 5 minutes.

### Quiescent cells preparation

To make nitrogen starvation induced quiescent fission yeast cells (Q_-N_ cells), a proper volume of proliferating culture (OD_600_ = 0.5 −1.0) was harvested by centrifugation and washed three times in EMM-N-C liquid, then re-suspended into 200ml EMM-N and its variants to the concentration of designed initial OD (OD_600_ = 0.15, 0.3, 0.4, 0.5, 0.6, 0.8, 1.0). The time when all the washes finished was recorded as the T0 for nitrogen starvation. For preparing stationary cells (ST cells), 200ml growth culture was split out from the remaining proliferating culture and continued incubating at 30°C along with the nitrogen starvation samples. The T0 for Q_-N_ conditions were arbitrarily assigned to ST condition. During continuous starvation, 1ml culture was taken out at each indicated time-point and split into 3 parts, 200μl for serial dilution for CFU plating, 200μl for agarose dish sampling, remaining 600μl for supernatant collection and glucose measurement. The supernatant was stored at −20°C before batch measuring. The sampling and measurements were continued for one month. All liquid yeast cultures were incubated at 30°C with shaking (220 rpm).

### Time-lapse imaging

Time-lapse images were acquired at 5minutes interval with a Nikon Eclipse Ti Microscope equipped with an automated stage, a perfect-focus-system, an 60x oil immersion objective, an Evolve EMCCD camera, and built in NIS-Elements softerware. For imaging proliferating cells, agarose dish loaded with cells were pre-cultured for 2hours on microscope at 30°C and then imaged for 12hours. For imaging quiescent cells, time-lapse program was immediately run after imaging program settle down. Before time-lapse imaging, one loop capture was performed to verify the program works correctly and the lens oil disperses homogenously. This one-loop images were subjected for cell segmentation and cell size extraction with the CellSeg software^52^. After started the time-lapse imaging program, recorded the time as T0’. The imaging duration was set as 16 hours at day1 and extended accordingly in the following days. During imaging, if cells were found off focus, the imaging program was paused to re-adjust the focus. The pause interval was automatically recoded by the program and taken into account for calculating the single cell recovery time. All microscope agarose dishes were maintained at 30°C during imaging.

### Yeast cells immobilization

Before fluorescence imaging, yeast cells were immobilized to the microscope plate which was coated with Concanavalin A (Sigma-Aldrich, Cat# C2010). For coating 384well microscope plate (Cellvis, Cat# P384-1.5H-N), fill each well with 20μl Concanavalin A water solution (2mg/ml) to cover the glass surface. After 2mins’ coating, remove the solution and dry the plate for several hours. To immobilize yeast cells, drop 5μl cell suspension into the coated well, then add 15μl 1x PBS into the well to achieve a proper cell density. Centrifuge the plate for 30 seconds at 500g to sediment and immobilize yeast cells.

### BODIPY493/503 staining

BODIPY493/503 (Invitrogen, Cat# D3922) was dissolved in DMSO at 1mg/ml and then divided into 100μl aliquots and stored at −20°C under dark as 1000x stock. For lipids staining, add 1μl 1mg/ml BODIPY493/503 stock into 1ml yeast suspension and stain for 10min under dark. After staining, load 5μl of each stained sample into the wells of 384-well microscope plate which coated with Concanavalin A and then add 15μl 1x PBS into each sample and mix well. Centrifuge the plate for 30 seconds at 500g to sediment and immobilize yeast cells. After immobilization, detect BODIPY493/503 fluorescence with a confocal microscope. If not specially mentioned, the stained samples were not washed by 1x PBS before loading into wells.

### Confocal fluorescence microscope

To investigate the organelle morphology and target gene expression under quiescence state, double-color fluorescence strains were imaged with confocal fluorescence microscope. Images were taken with a 100x oil TIRF objective plus 1.5x intermediate magnification. During imaging, yeast cells were sliced for 15 layers within 4um thickness. More than 100 single cells were imaged for each sample.

### CFU plating

Before plating, cultures were serially diluted to appropriate concentration (100-150 cells per 100μl) by EMM liquid. The diluted concentrations for samples with different initial OD were 7.0×1^-5^, 3.5×10^-5^, 2.5×10^-5^, 2.0×10^-5^, 1.5×10^-5^, 1.25×10^-5^, 1.0×10^-5^, and 1.0×10^-5^ for initial OD of 0.15, 0.3, 0.4, 0.5, 0.6, 0.8, 1.0 and ST respectively. Then 200μl end dilution was plated on to each YE agar plate with 4 repeats plated for each sample. After 3 days of incubation at 30°C, the colonies on each plate were counted. The survival rate measured by CFU method was calculated as the percentage of colony counts at each time-point relative to the reference counts at day 1.

### Glucose concentration measurement

The culture medium glucose concentration was measured by Glucose Oxidase Method with a commercial assay kit (Applygen Technologies, Cat# E1010). Samples were the −20°C stocks of supernatant collected during each imaging experiment.

### Image analysis

Time-lapse images were exported with NIS-elements software. Images were analyzed with the software “Cellseg”^52^. All cells in the analyzed image were segmented. Those edge cells which later moved out of the field during time-lapse imaging were excluded for statistical analysis. More than 100 single cells were analyzed for each imaged sample (cell number details see Supplementary Tables S1–S3).

For single cell recovery time (T_recovery_) measurement, the single cell recovery behavior of quiescent cell was traced by eye and the image frame number “m” of first cell division was recorded by hand. The T_recovery_ for each single cell was defined as the time duration starting from dropping cells on agarose dish until the first cell division finished. T_recovery_ was the imaging time obtained from imaging data plus experimental time. Generally, T_recovery_ (minute) = 5(m-1) +T0’-T0, with T0 and T0’ are hand-recorded time during microscope agarose dish preparation and time-lapse imaging (see Time-lapse imaging). If there is a pause during imaging due to refocus, the pause time is taken into account. The survival rate measured by imaging was calculated as the percentage of fully recovered cells which finished first cell division in the analyzed population.

For fluorescence image analysis, all slices for each fluorescence channel were stacked into one gray image by Z projection with maximum tensity with the software Fiji^53^. The Z-projected gray image of each channel was background-subtracted and then merged into one RGB image. The “substract background” function of Fiji was used and the rolling ball diameter was set as 30.

For cell size measurement, images were analysis with the software “Cellseg”^52^. All cells fully presented in the image filed were segmented. All imprecise segmentations were corrected by hand using the build-in function of “edit segmentation” of the software “Cellseg”. The measurements of cell area and cellular fluorescence were exported by the “export” tool in the Cellseg.

### cDNA library preparation and RNA-seq

RNA was isolated using TRIzol (Thermo Fisher Scientific, Waltham, MA, USA). mRNA molecules were purified using the poly-T oligo attached magnetic beads following which the mRNA was fragmented and primed for cDNA synthesis. cDNA libraries were pair-end 150-bp sequenced on the DNBseq platform at BGI (Wu Han, China).

### RNA-seq data preprocessing

Raw sequences were filtered by SOAPnuke^54^ HISAT2^55^ was used to align the RNA sequencing reads to the Schizosaccharomyces pombe reference genome (https://www.pombase.org/downloads/genome-datasets). Paired-end cleaned reads were mapped to the reference genome using Bowtie2^56^. The number of mapped reads covering each gene and FPKM were calculated using RSEM^57^

### RNA-seq data analysis

To study the similarities between all samples, we performed log2 transformation on the FPKM matrix and calculated the Pearson correlation coefficients between the two samples. The log2 transformed expression matrix was subjected to Principle Component Analysis to visualize the sequential transition of transcriptome.

For gene expression clustering analysis, FPKM was log2 transformed. The R package WGCNA^58^ was used to detect clusters of highly correlated genes. Clustering results were visualized as heat maps using the R package pheatmap. The GO biological process slim table from the Pombe website (https://www.pombase.org/browse-curation/fission-yeast-bp-go-slimterms) was used as an annotation for the genes. GO terms enriched in WGCNA clusters were detected using the function ‘enricher’ in the R package clusterProfiler^59^. The average expression levels of genes included in the GO terms with Holm-Bonferroni-corrected p-value < 0.05 were shown in Figure 5B.

To globally visualize the functional modules regulated differently under different cell densities and glucose concentrations during the quiescence development and cellular ageing, the functional enrichment of all samples relative to proliferative state samples was shown in Figure 5C. For this analysis, differentially expressed genes between all samples and proliferative state samples were detected using the R package DESeq2^60^. The cut-off for significance was the adjusted p-value <0.05 and the fold difference >2. This analysis generated a list of significantly up and down-regulated genes for each group. These gene lists were analyzed using the function ‘enricher’ in the R package clusterProfiler^59^. GO slims with a Holm-Bonferroni-corrected p-value < 0.05 were considered to be significantly enriched within the lists of genes. GO categories enriched less than 2 out of the 44 groups were discarded. Fold enrichment of 27 significantly enriched GO slims was shown in heatmap.

## Supporting information

data sheets

## SUPPLEMENTAL INFORMATION

Supplemental information includes six figures, seven tables, and five movies.

## MATERIALS AVAILABILITY

All unique reagents generated in this study are available from the Lead Contact without restriction.

## DATA AND CODE AVAILABILITY

Single cell data of recovery time and cell size and the RNA-seq data of gene expression FPKM value generated during this study are provided as datasheet. The original sequence data is available in NCBI BioProject database with the ID number of PRJNA900375. This study did not generate any unique code.

## ACKNOWLEDGMENTS

We thank all members of the Tang Laboratory for discussion and support. We also thank Dr. Dao-chun Kong (Peking University), Dr. Qi OUYANG (Peking University), Dr. Fang-ting Li (Peking University), Dr. Qing Li (Peking University), Dr. Li-lin Du (NIBS), Dr. Meng-Qiu Dong (NIBS), Dr. Hui-qiang Lou (China Agricultural University), and Dr. Dong-gen Luo (Peking University) for discussion. This work was supported by the National Key Research and Development Program of China (2018YFA0900700, 2021YFF1200500) and National Science Foundation of China (NSFC 12090053 and NSFC 32088101).

## AUTHOR CONTRIBUTIONS

Q.L. and C.T. conceived the project; Q.L. designed and performed all the experiments, analyzed most image data; N.S., Q. L., and X.Y. analyzed RNA-seq data; Z.Z., C.H., Z.Y., and H.S. also contributed to image data analysis; C.T., J.C., and X.Y. supervised the project; Q.L., N. S., X.Y., and C.T. wrote the paper.

---

**Supplementary Table S1.**
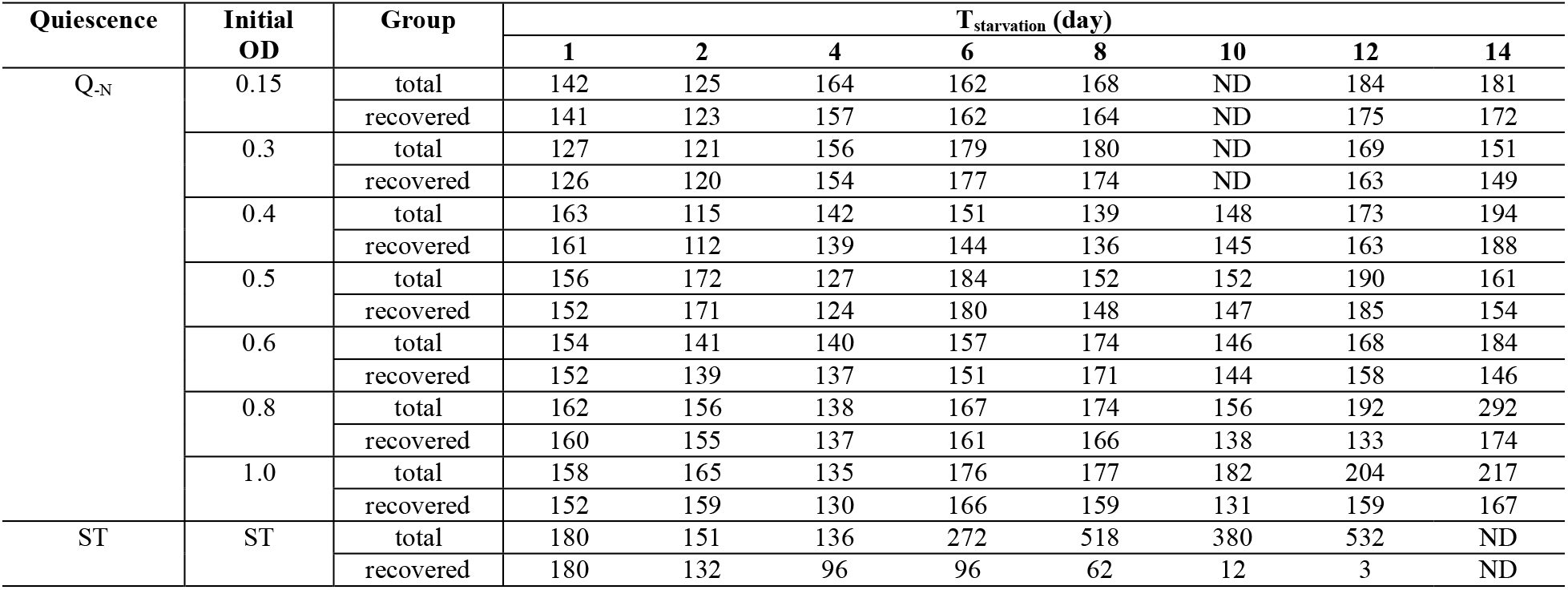
Analyzed numbers of single quiescent cells with various initial OD from imaging data. Q_-N_: nitrogen-starvation induced quiescence, ST: stationary phase, ND: no data

**Supplementary Table S2.**
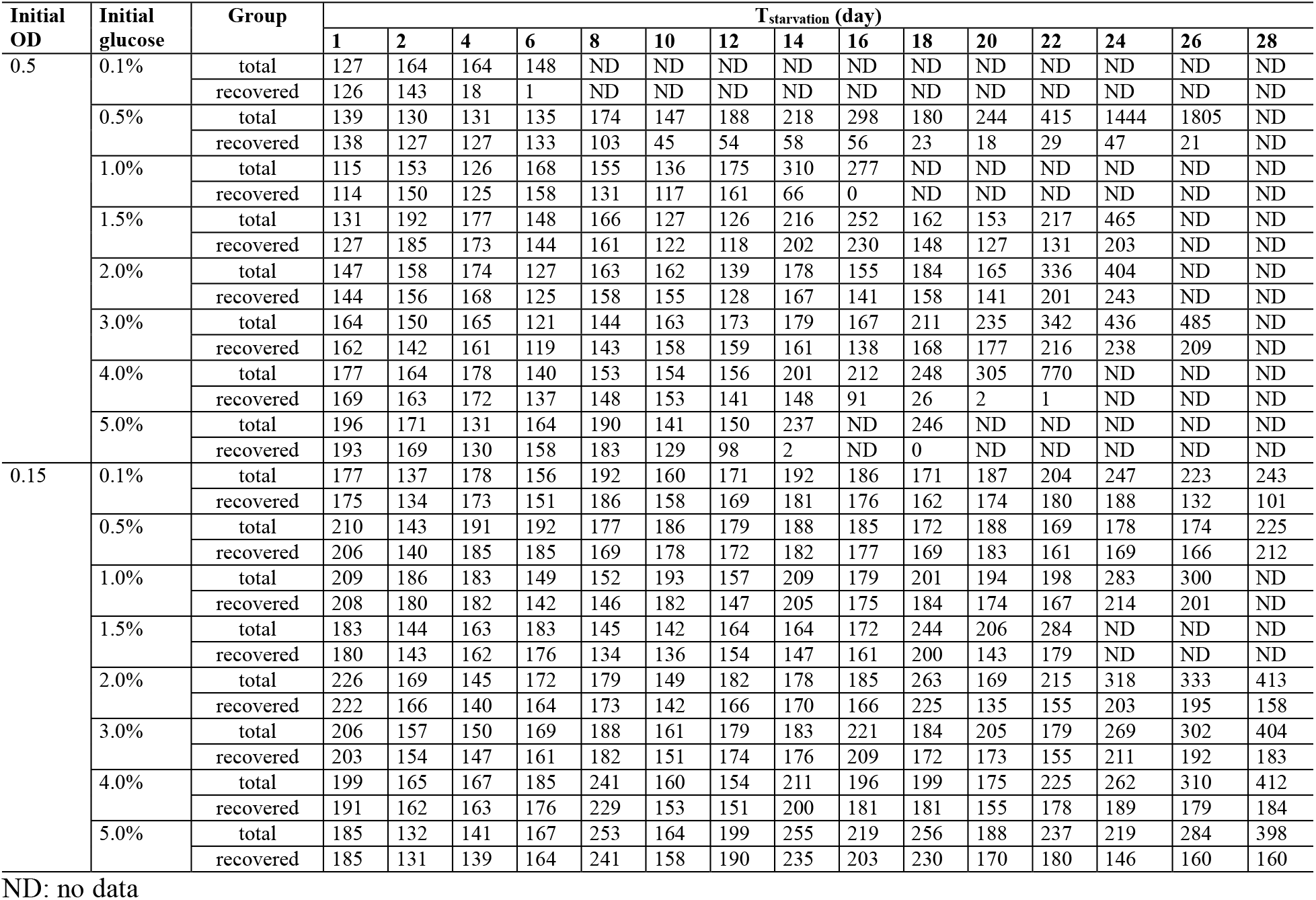
Analyzed numbers of single Q_-N_ cells with various initial glucose from imaging data.

| Initial OD | Initial glucose | Group | T<sub>starvation</sub> (day) |  |  |  |  |  |  |  |  |  |  |  |  |  |  |
| --- | --- | --- | --- | --- | --- | --- | --- | --- | --- | --- | --- | --- | --- | --- | --- | --- | --- |
|  |  |  | 1 | 2 | 4 | 6 | 8 | 10 | 12 | 14 | 16 | 18 | 20 | 22 | 24 | 26 | 28 |
| 0.5 | 0.1% | total | 127 | 164 | 164 | 148 | ND | ND | ND | ND | ND | ND | ND | ND | ND | ND | ND |
|  |  | recovered | 126 | 143 | 18 | 1 | ND | ND | ND | ND | ND | ND | ND | ND | ND | ND | ND |
|  | 0.5% | total | 139 | 130 | 131 | 135 | 174 | 147 | 188 | 218 | 298 | 180 | 244 | 415 | 1444 | 1805 | ND |
|  |  | recovered | 138 | 127 | 127 | 133 | 103 | 45 | 54 | 58 | 56 | 23 | 18 | 29 | 47 | 21 | ND |
|  | 1.0% | total | 115 | 153 | 126 | 168 | 155 | 136 | 175 | 310 | 277 | ND | ND | ND | ND | ND | ND |
|  |  | recovered | 114 | 150 | 125 | 158 | 131 | 117 | 161 | 66 | 0 | ND | ND | ND | ND | ND | ND |
|  | 1.5% | total | 131 | 192 | 177 | 148 | 166 | 127 | 126 | 216 | 252 | 162 | 153 | 217 | 465 | ND | ND |
|  |  | recovered | 127 | 185 | 173 | 144 | 161 | 122 | 118 | 202 | 230 | 148 | 127 | 131 | 203 | ND | ND |
|  | 2.0% | total | 147 | 158 | 174 | 127 | 163 | 162 | 139 | 178 | 155 | 184 | 165 | 336 | 404 | ND | ND |
|  |  | recovered | 144 | 156 | 168 | 125 | 158 | 155 | 128 | 167 | 141 | 158 | 141 | 201 | 243 | ND | ND |
|  | 3.0% | total | 164 | 150 | 165 | 121 | 144 | 163 | 173 | 179 | 167 | 211 | 235 | 342 | 436 | 485 | ND |
|  |  | recovered | 162 | 142 | 161 | 119 | 143 | 158 | 159 | 161 | 138 | 168 | 177 | 216 | 238 | 209 | ND |
|  | 4.0% | total | 177 | 164 | 178 | 140 | 153 | 154 | 156 | 201 | 212 | 248 | 305 | 770 | ND | ND | ND |
|  |  | recovered | 169 | 163 | 172 | 137 | 148 | 153 | 141 | 148 | 91 | 26 | 2 | 1 | ND | ND | ND |
|  | 5.0% | total | 196 | 171 | 131 | 164 | 190 | 141 | 150 | 237 | ND | 246 | ND | ND | ND | ND | ND |
|  |  | recovered | 193 | 169 | 130 | 158 | 183 | 129 | 98 | 2 | ND | 0 | ND | ND | ND | ND | ND |
| 0.15 | 0.1% | total | 177 | 137 | 178 | 156 | 192 | 160 | 171 | 192 | 186 | 171 | 187 | 204 | 247 | 223 | 243 |
|  |  | recovered | 175 | 134 | 173 | 151 | 186 | 158 | 169 | 181 | 176 | 162 | 174 | 180 | 188 | 132 | 101 |
|  | 0.5% | total | 210 | 143 | 191 | 192 | 177 | 186 | 179 | 188 | 185 | 172 | 188 | 169 | 178 | 174 | 225 |
|  |  | recovered | 206 | 140 | 185 | 185 | 169 | 178 | 172 | 182 | 177 | 169 | 183 | 161 | 169 | 166 | 212 |
|  | 1.0% | total | 209 | 186 | 183 | 149 | 152 | 193 | 157 | 209 | 179 | 201 | 194 | 198 | 283 | 300 | ND |
|  |  | recovered | 208 | 180 | 182 | 142 | 146 | 182 | 147 | 205 | 175 | 184 | 174 | 167 | 214 | 201 | ND |
|  | 1.5% | total | 183 | 144 | 163 | 183 | 145 | 142 | 164 | 164 | 172 | 244 | 206 | 284 | ND | ND | ND |
|  |  | recovered | 180 | 143 | 162 | 176 | 134 | 136 | 154 | 147 | 161 | 200 | 143 | 179 | ND | ND | ND |
|  | 2.0% | total | 226 | 169 | 145 | 172 | 179 | 149 | 182 | 178 | 185 | 263 | 169 | 215 | 318 | 333 | 413 |
|  |  | recovered | 222 | 166 | 140 | 164 | 173 | 142 | 166 | 170 | 166 | 225 | 135 | 155 | 203 | 195 | 158 |
|  | 3.0% | total | 206 | 157 | 150 | 169 | 188 | 161 | 179 | 183 | 221 | 184 | 205 | 179 | 269 | 302 | 404 |
|  |  | recovered | 203 | 154 | 147 | 161 | 182 | 151 | 174 | 176 | 209 | 172 | 173 | 155 | 211 | 192 | 183 |
|  | 4.0% | total | 199 | 165 | 167 | 185 | 241 | 160 | 154 | 211 | 196 | 199 | 175 | 225 | 262 | 310 | 412 |
|  |  | recovered | 191 | 162 | 163 | 176 | 229 | 153 | 151 | 200 | 181 | 181 | 155 | 178 | 189 | 179 | 184 |
|  | 5.0% | total | 185 | 132 | 141 | 167 | 253 | 164 | 199 | 255 | 219 | 256 | 188 | 237 | 219 | 284 | 398 |
|  |  | recovered | 185 | 131 | 139 | 164 | 241 | 158 | 190 | 235 | 203 | 230 | 170 | 180 | 146 | 160 | 160 |

ND: no data

**Supplementary Table S3.**
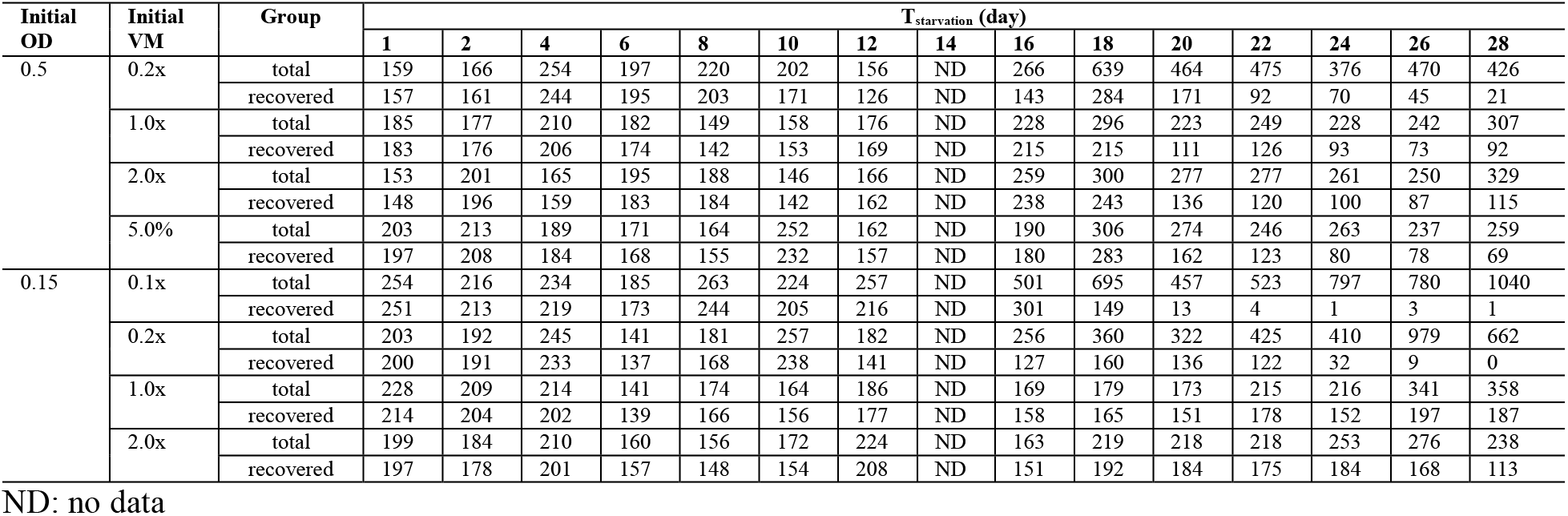
Analyzed numbers of single Q_-N_ cells with various initial VM from imaging data.

**Supplementary Table S4.**
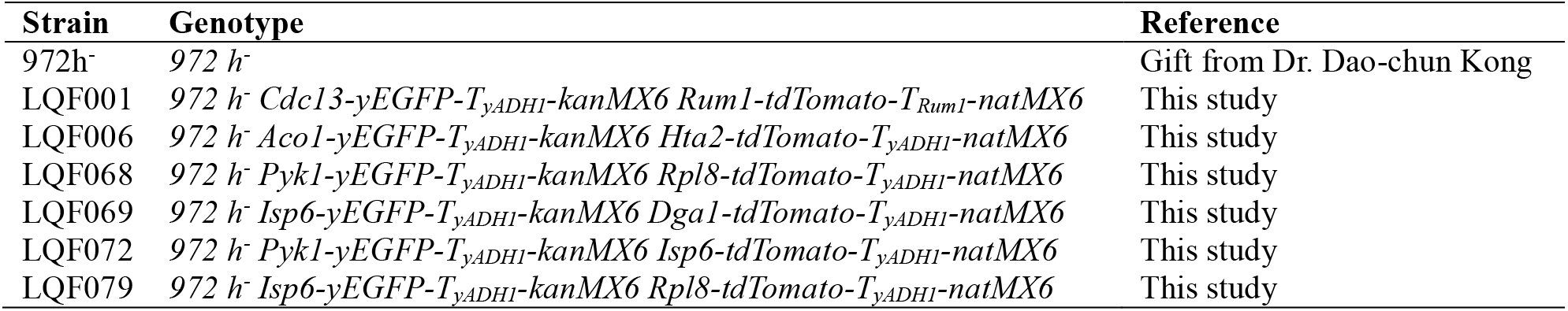
Fission yeast strains used in this study.

**Supplementary Table S5.**
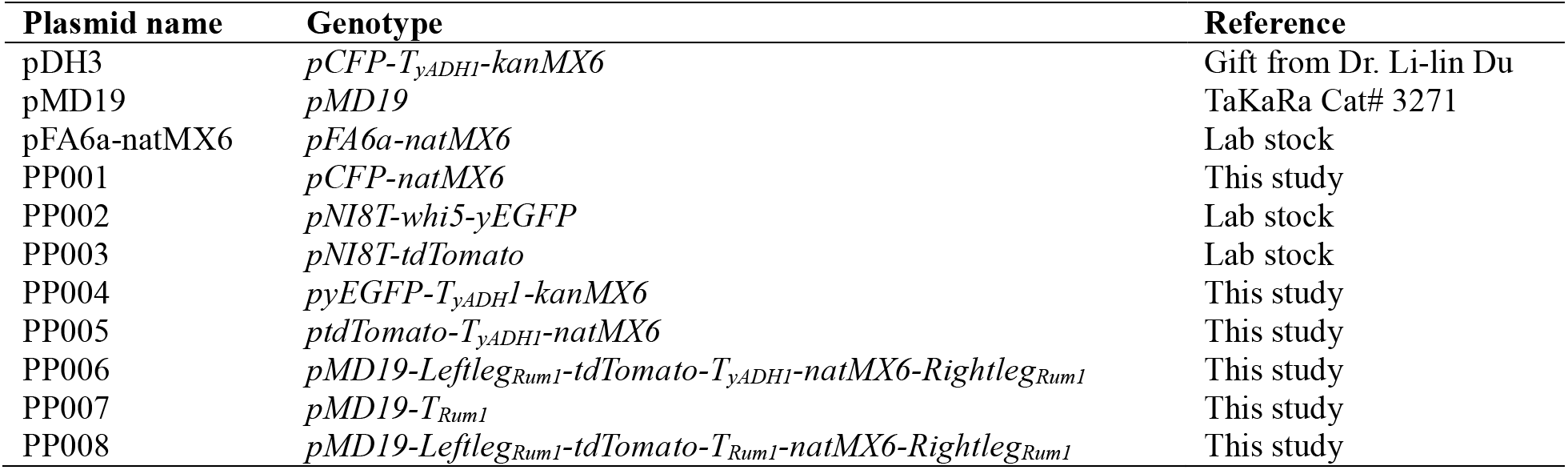
Plasmids used in this study.

**Supplementary Table S6.**
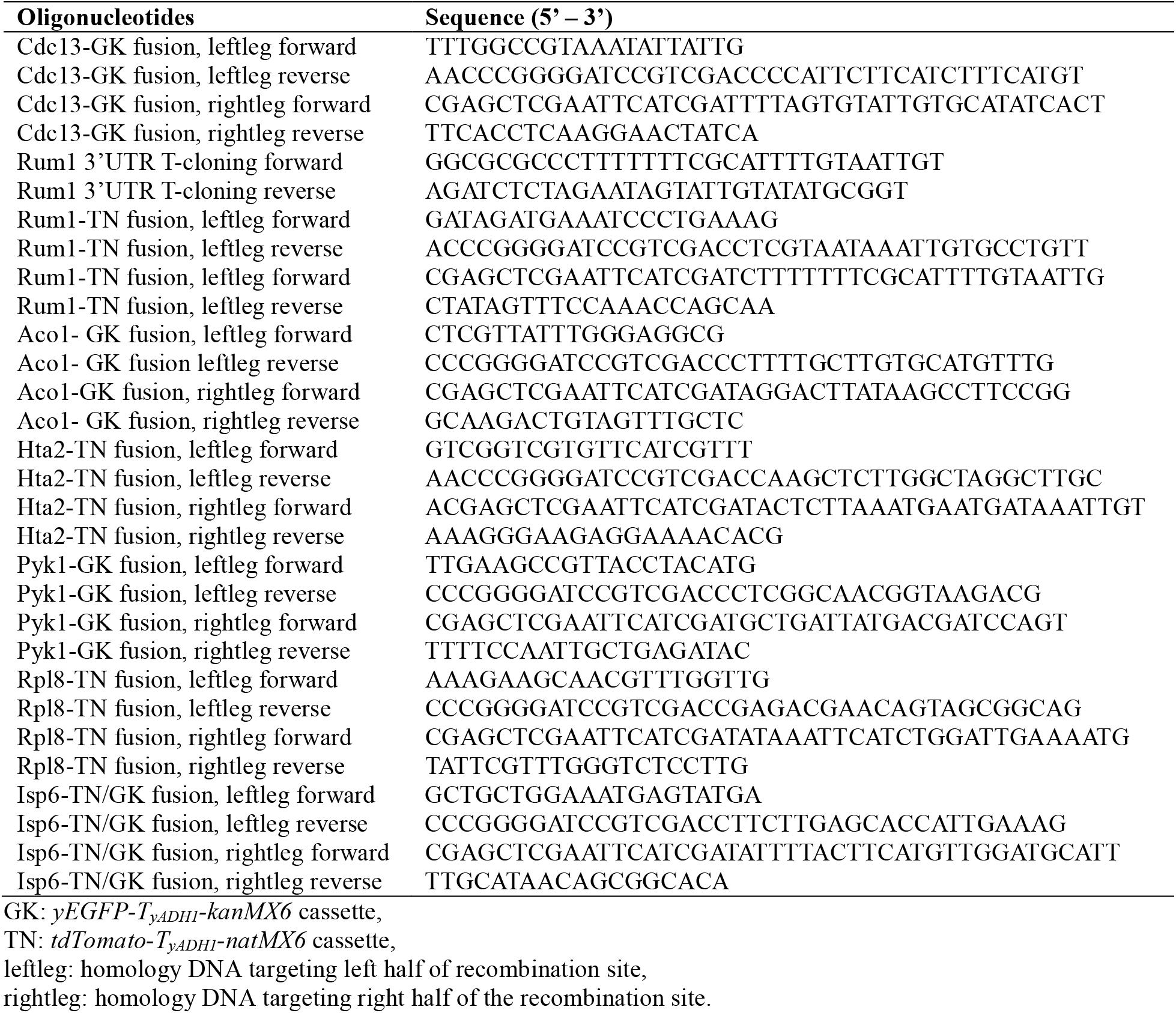
Oligonucleotides used in this study.

**Supplementary Table S7.**
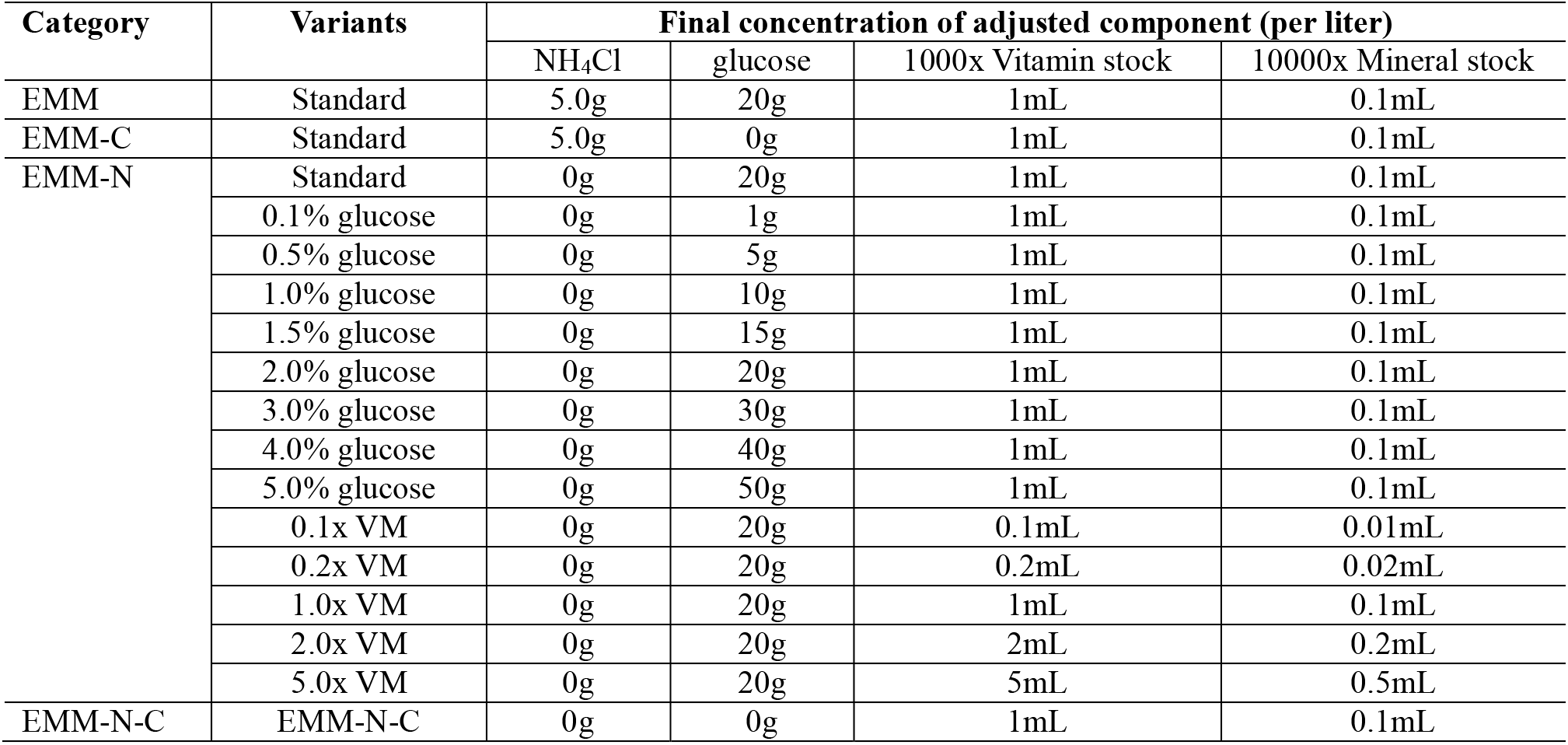
Synthetic media used in this study.

**Captions for Supplementary Movies S1 to S5**

**Movie S1.**

Nitrogen-starved quiescent cells accumulate LD during quiescence establishment.

**Movie S2.**

Proliferating cells store basal level LD.

**Movie S3.**

Recovery of Q_-N_ cells starved for 16 days under initial OD=0.5 with 0.5% initial glucose.

**Movie S4.**

Recovery of Q-N cells starved for 16 days under initial OD=0.5 with 2.0% initial glucose.

**Movie S5.**

Recovery of Q-N cells starved for 16 days under initial OD=0.5 with 4.0% initial glucose.

---

**Figure S1.**
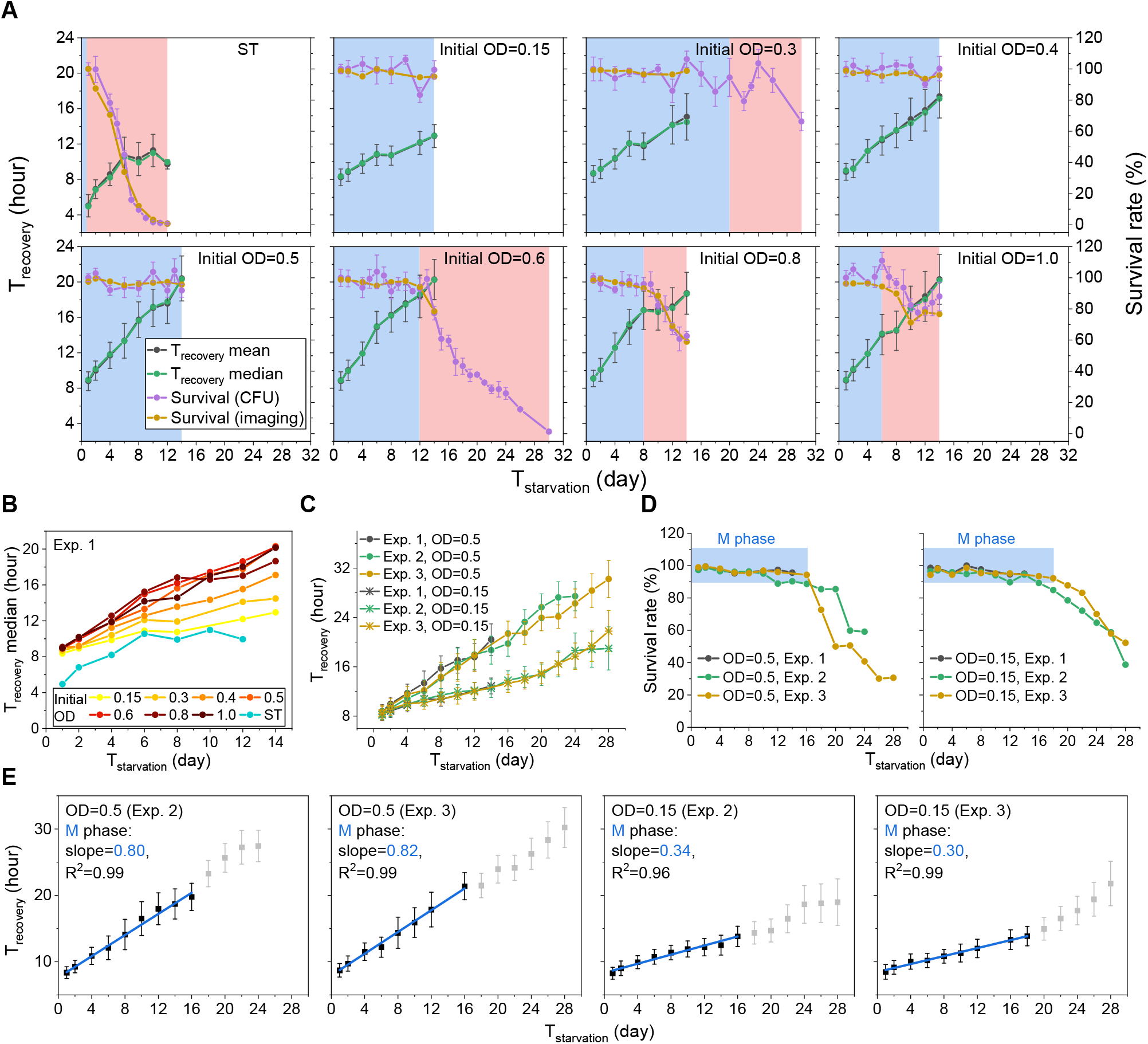
The linear aging behavior of quiescent cell is robust. **(A)** Multi plots of T_recovery_, survival rate, M phase and D phase under various OD conditions. Data for T_recovery_ mean (black) and survive rate by CFU (purple) are represented as mean ± SEM. M phase and D phase are indicated by the blue and red shadows respectively. Data condition is presented in the upright of each panel. **(B)** Comparison of T_recovery_ median under various OD conditions. (**C-D**) Comparison of T_recovery_ (**C**) and survival arte (**D**) measured in different experimental repeats under the initial OD of 0.5 and 0.15. Data for T_recovery_ in **C** are represented as mean ± SEM. (**E**) Linear fitting of T_recovery_ vs. T_starvation_ data during M phase for the experimental repeats. Data condition, linear fitting slope, and R^2^ are presented in the up left corner of each panel. Grayed dots represent D phase. Blue line represents fitting curve. Data are represented as mean ± SEM.

**Figure S2.**
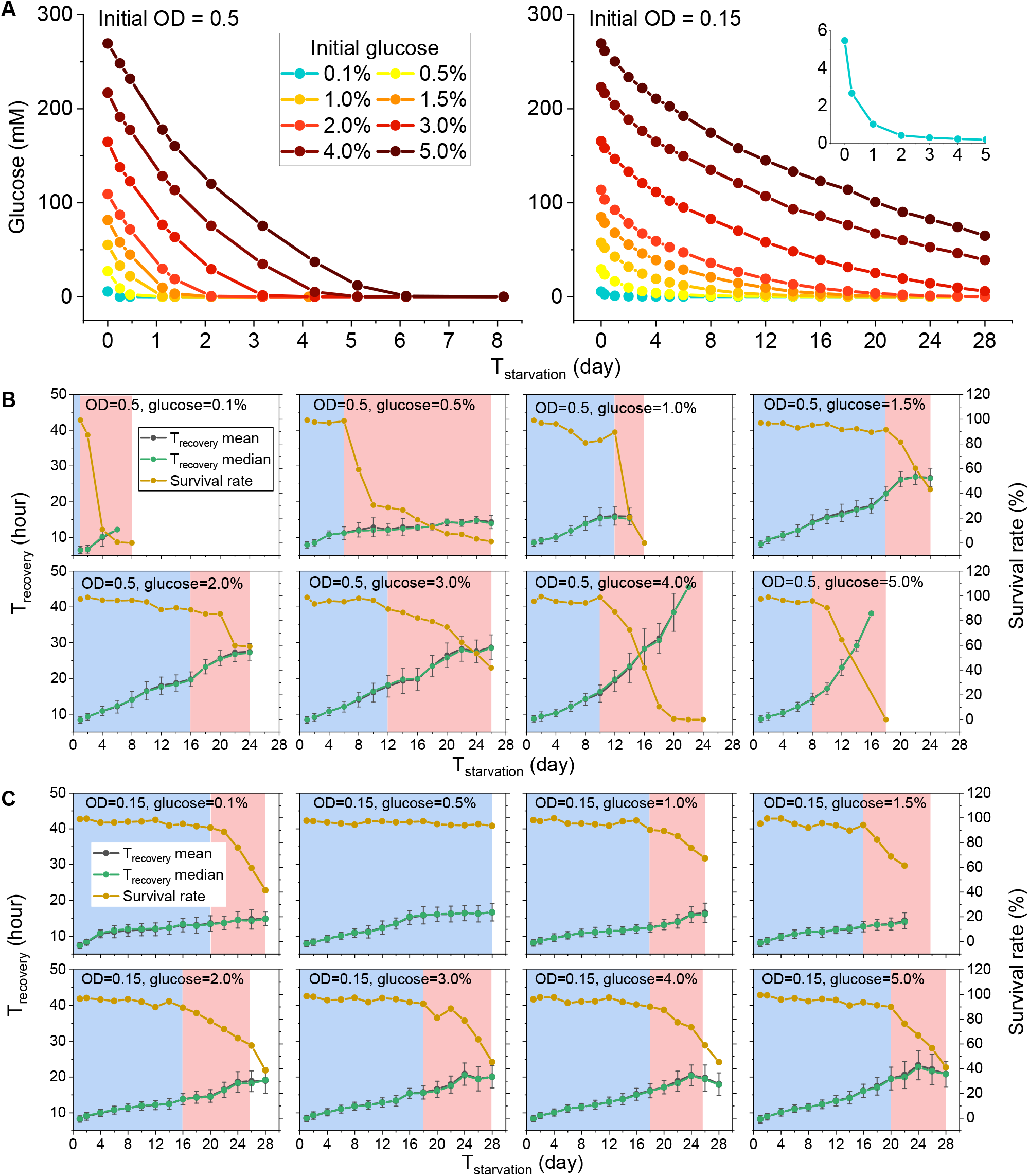
Measured T_recovery_, survival rate, M phase, and D phase under various glucose conditions under two representative ODs. **(A)** Glucose consuming curves. **(B-C)**, Multi plots of T_recovery_, survival rate, M phase and D phase under various glucose conditions with initial OD of 0.5 (**B**) and 0.15 (**C**). Data condition is presented in the top of each panel. M phase and D phase are indicated by the blue and red shadows respectively. Cell numbers for each data point are listed in Table S2.

**Supplementary Figure S3.**
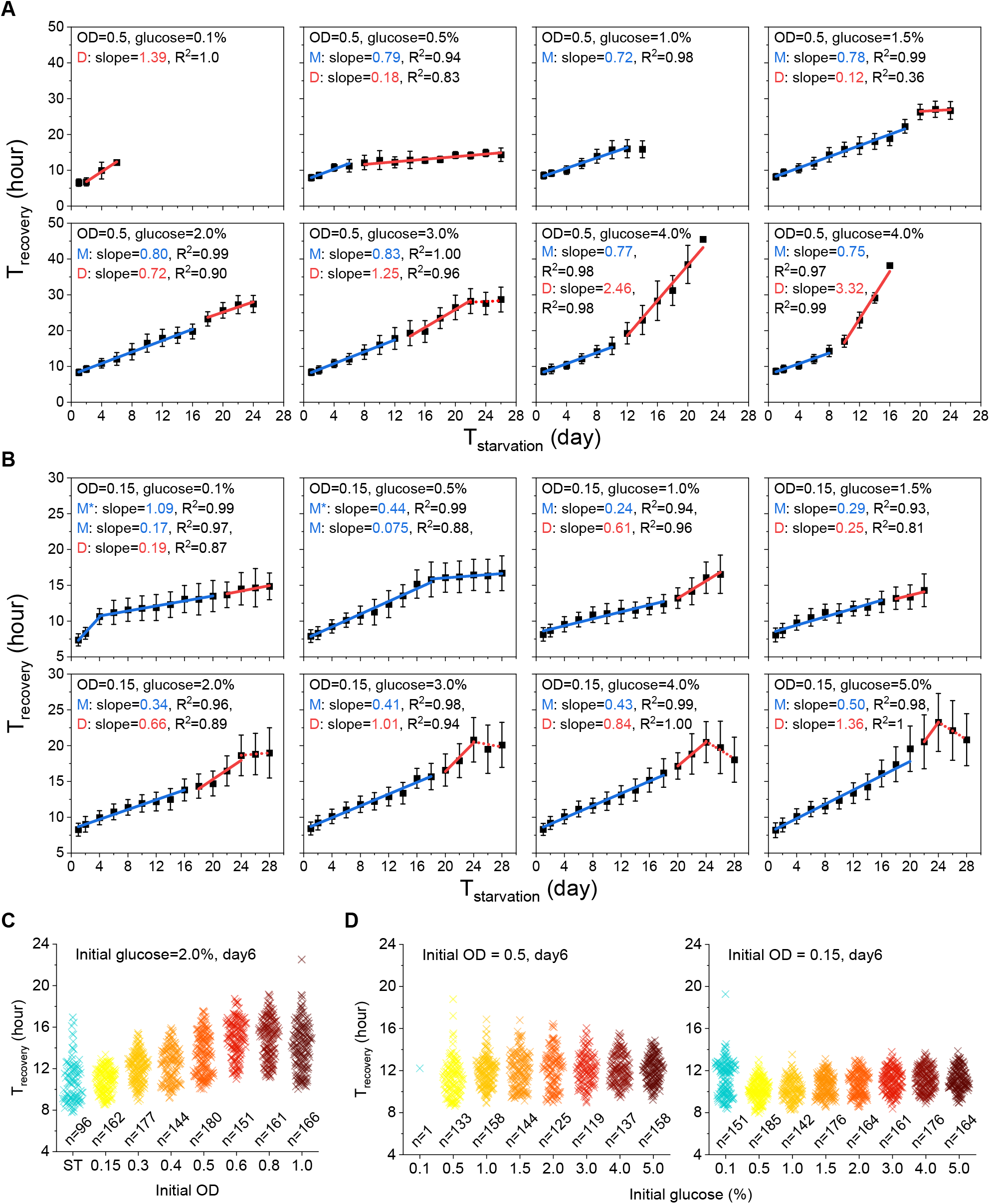
Comparison of T_recovery_ under various glucose conditions under two representative ODs. **(A-B), Linear fitting of T_recovery_ data during M phase (blue lines) and D phase (red lines) under various glucose conditions with the initial OD of 0.5 (A) and 0.15 (B). Data condition, linear fitting slope, and R^2^ are presented in the up left corner of each panel. Cell numbers for each data point are listed in Table S2.** **(C-D)** Comparison of T_recovery_ for quiescent cells at day6 under different OD (C) and different glucose concentrations with two representative ODs (D). Cell numbers are indicated.

**Figure S4.**
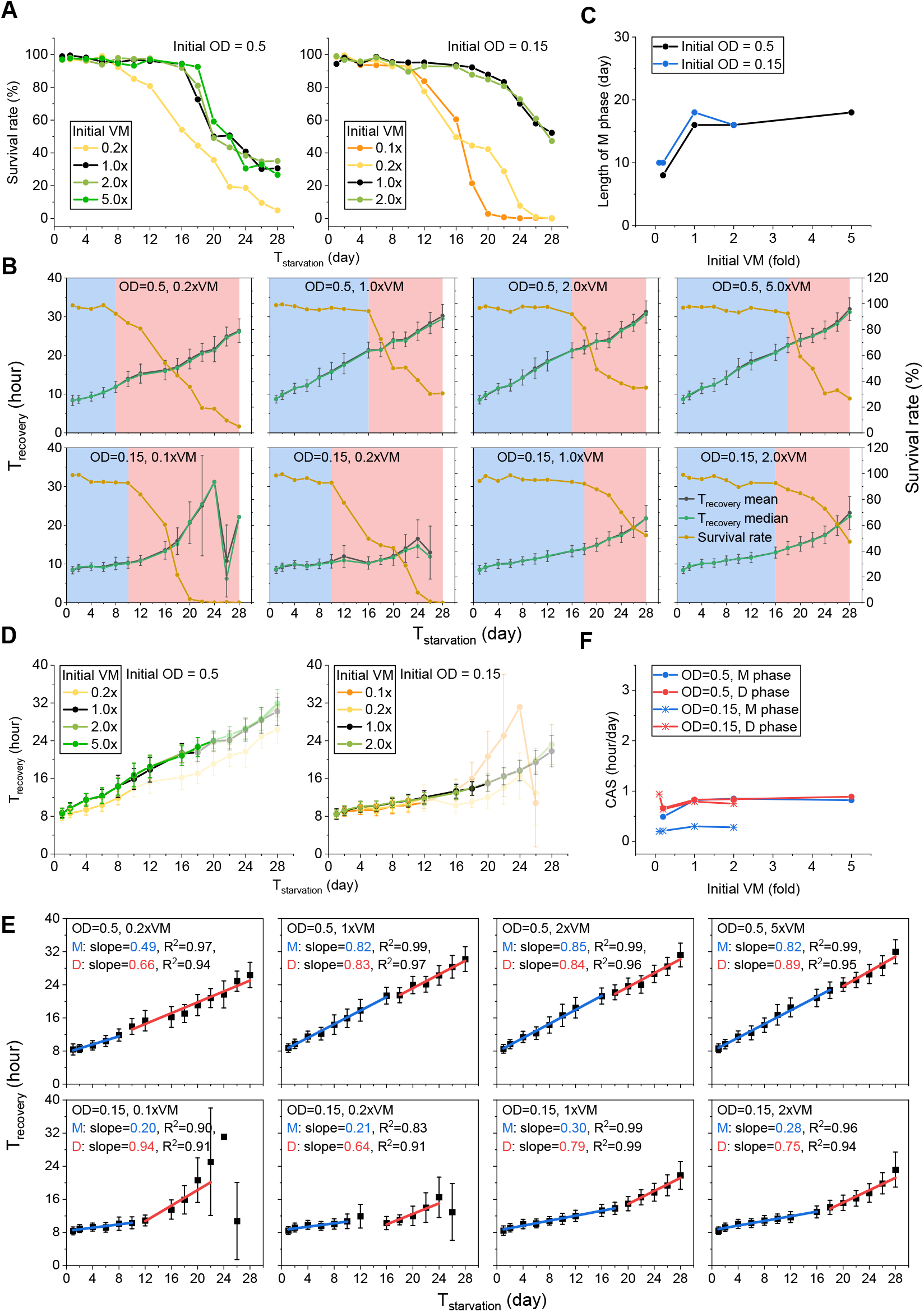
Single cell aging behaviors under various VM conditions under two representative ODs. **(A-D)** Measured survival rate (**A**), M phase and D phase (**B**), length of M phase (**C**), and T_recovery_ (**D**) under various <u>v</u>itamin and <u>m</u>ineral concentrations (VM) under two representative ODs. **(E)** Linear fitting of T_recovery_ data in **D** for each condition during M phase (blue lines) and D phase (red lines). Data condition, linear fitting slope, and R^2^ are presented in the up left corner of each panel. (**F**) Comparison of CAS measured in **E**. Cell numbers for each data point are listed in Table S3.

**Figure S5.**
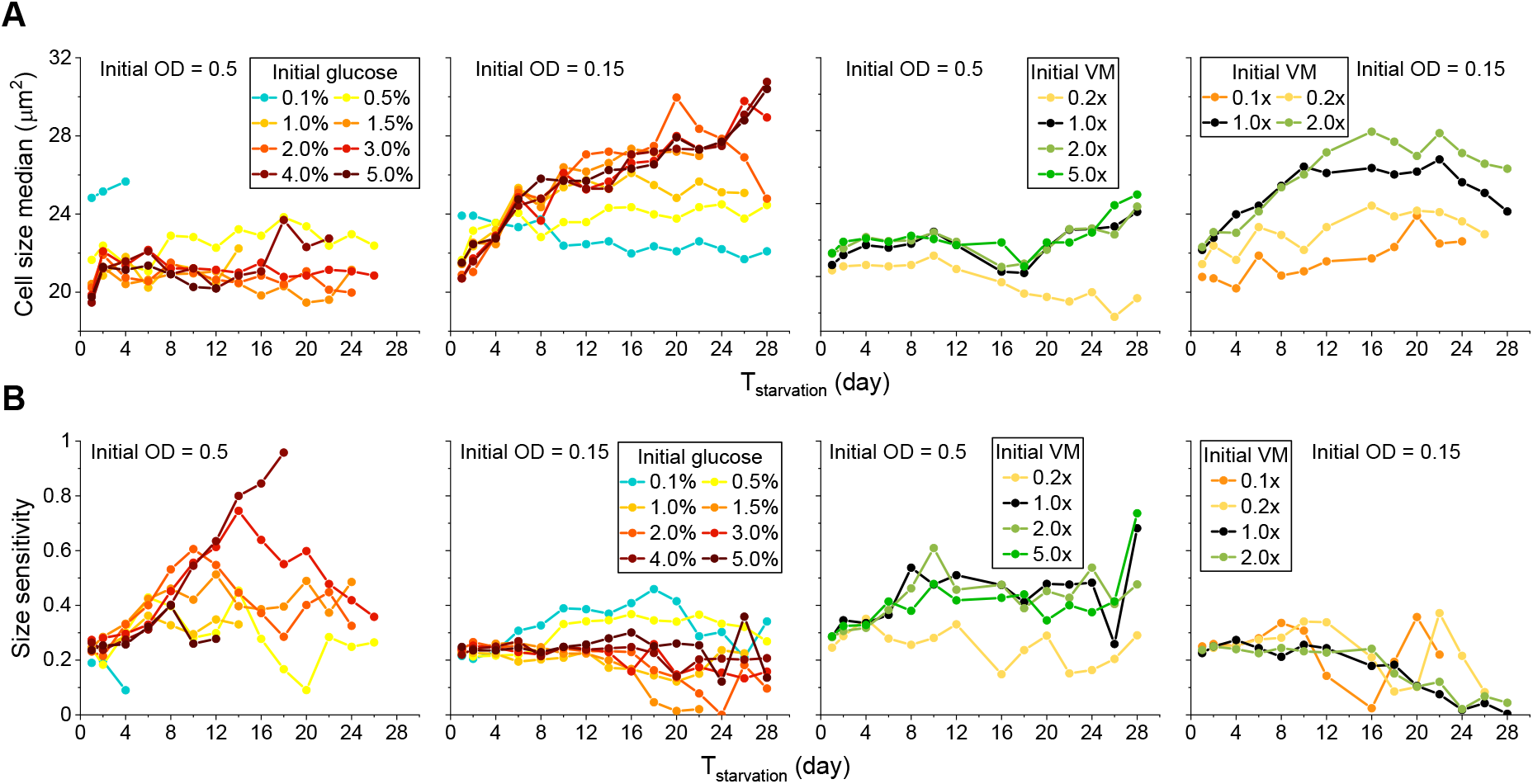
Comparison of quiescent cell size and size sensitivity under various glucose and VM conditions. **(A-B)** The change of quiescent cell size (**A**) and size sensitivity (**B**) of Q_-N_ cells under various glucose and VM conditions under the initial OD of 0.5 and 0.15. Cell size in **A** is presented by the median value and data condition is presented in each panel. Cell numbers are listed in Table S2 and S3.

**Figure S6.**
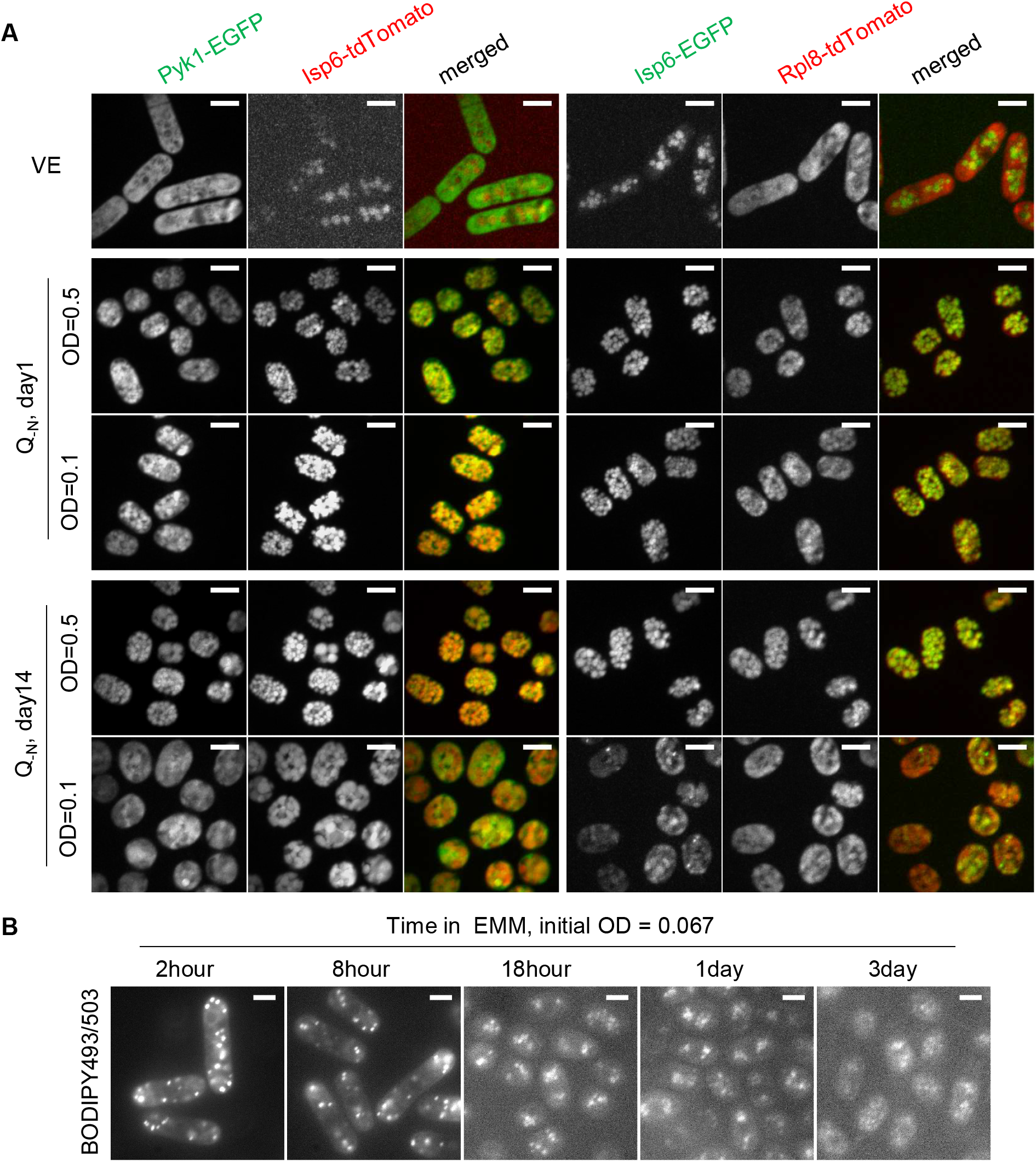
The co-localization of glycolysis enzyme and ribosome with vacuolar, decrease of lipids accumulation at stationary phase. **(A)** The localization of Pyk1-EGFP, Isp6-tdTomato, Isp6-EGFP, and Rpl8-tdTomato in day1 and day14 Q_-N_ cells under the initial OD of 0.5 and 0.1. **(B)** Change of lipids accumulation during ST cell ageing. Lipids are stained with BODIPY493/503. The scale bar is 5μm.

