## supplemental figures S1-S6 for "Initial niche condition determines the aging speed and regenerative activity of quiescent cells"

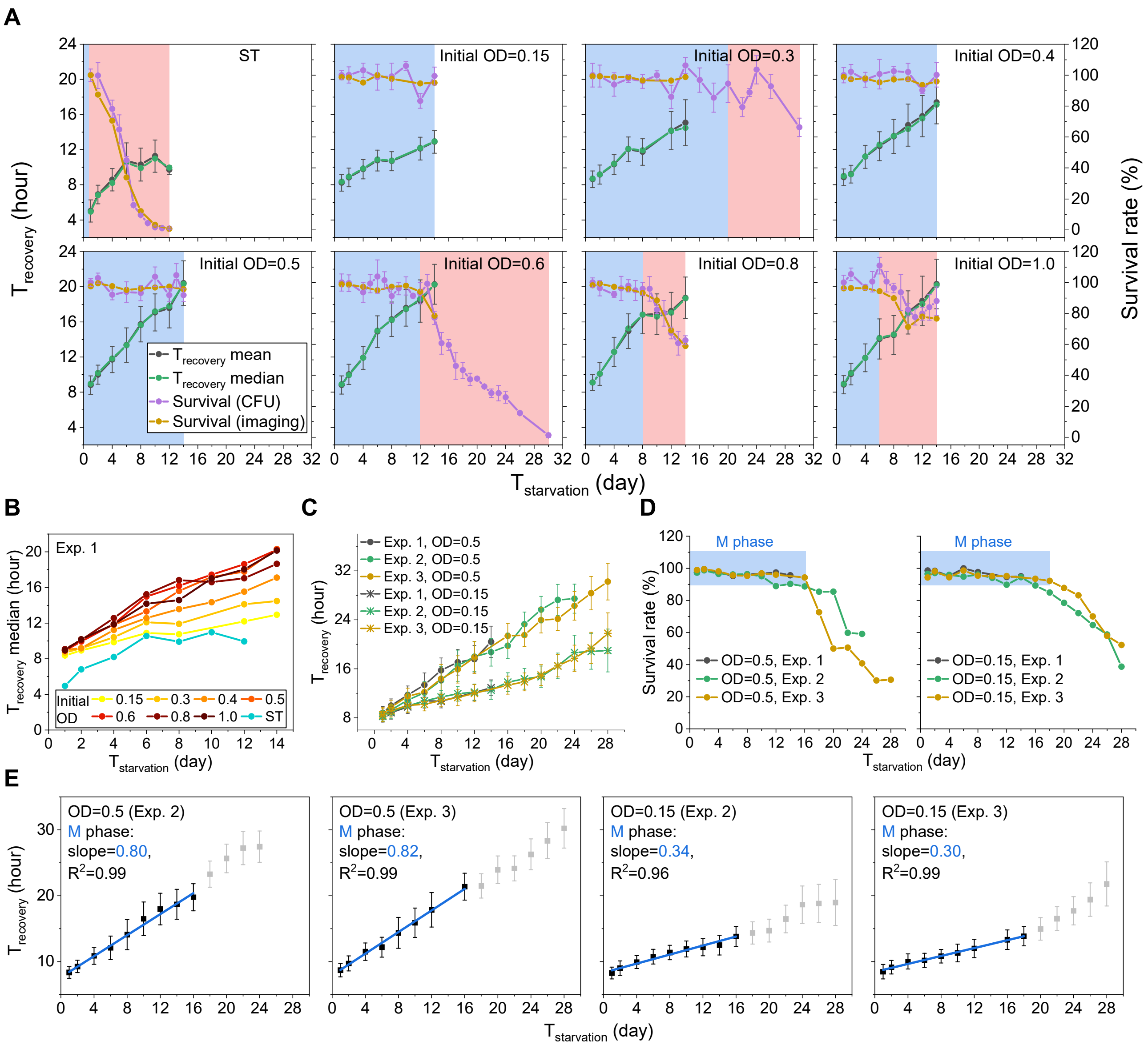


**Figure S1. The linear aging behavior of quiescent cell is robust.**

**(A)** Multi plots of $T_{recovery}$, survival rate, M phase and D phase under various OD conditions. Data for $T_{recovery}$ mean (black) and survive rate by CFU (purple) are represented as mean ± SEM. M phase and D phase are indicated by the blue and red shadows respectively. Data condition is presented in the upright of each panel.

**(B)** Comparison of $T_{recovery}$ median under various OD conditions.

(**C**-**D**) Comparison of $T_{recovery}$ (**C**) and survival arte (**D**) measured in different experimental repeats under the initial OD of 0.5 and 0.15. Data for $T_{recovery}$ in **C** are represented as mean ± SEM.

(**E**) Linear fitting of $T_{recovery}$ vs. $T_{starvation}$ data during M phase for the experimental repeats. Data condition, linear fitting slope, and $R^2$ are presented in the up left corner of each panel. Grayed dots represent D phase. Blue line represents fitting curve. Data are represented as mean ± SEM.

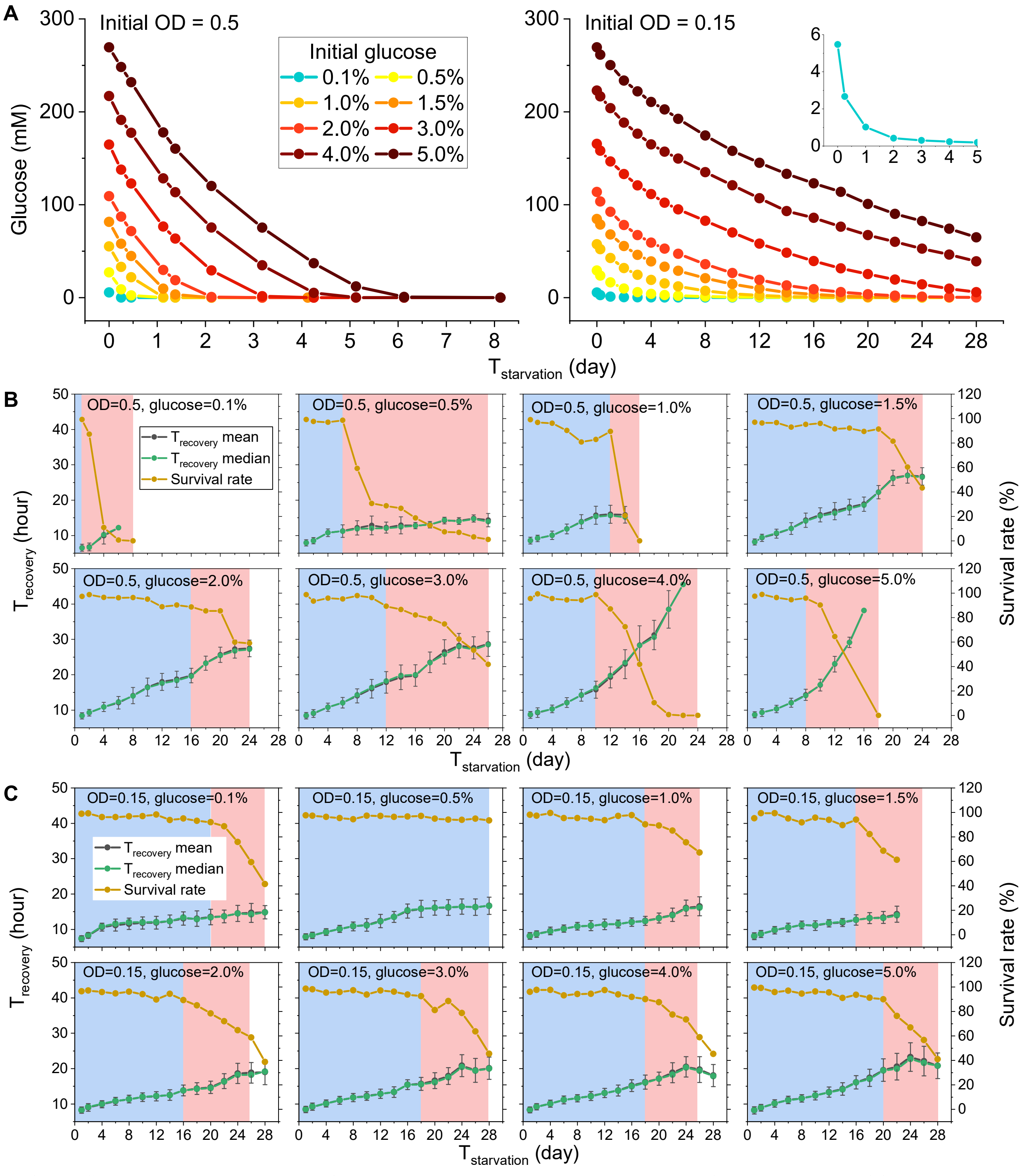


**Figure S2. Measured $T_{recovery}$, survival rate, M phase, and D phase under various glucose conditions under two representative ODs.**

**(A)** Glucose consuming curves.

**(B-C**), Multi plots of $T_{recovery}$, survival rate, M phase and D phase under various glucose conditions with initial OD of 0.5 (**B**) and 0.15 (**C**). Data condition is presented in the top of each panel. M phase and D phase are indicated by the blue and red shadows respectively. Cell numbers for each data point are listed in Table S2.

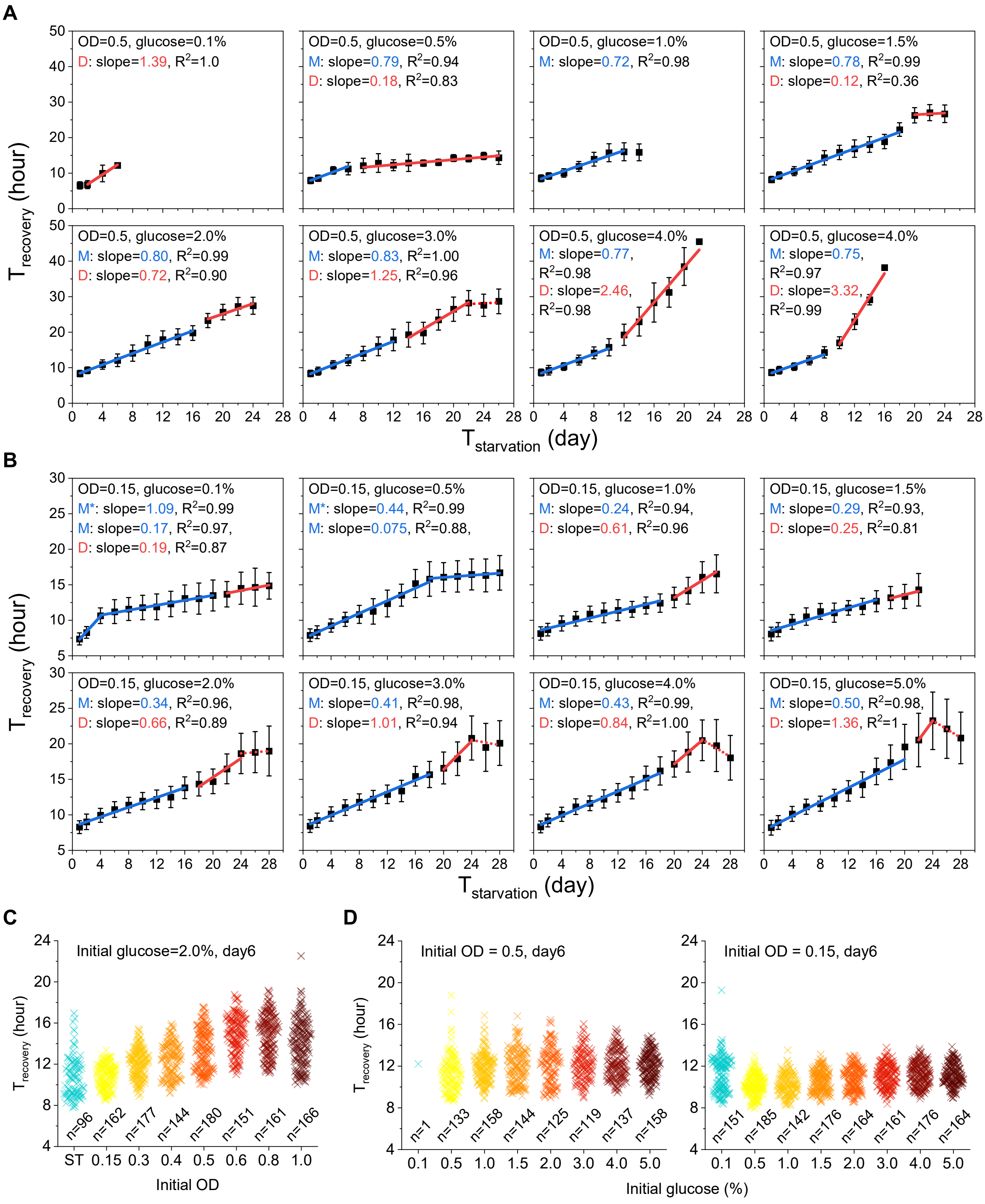


**Supplementary Figure S3. Comparison of $T_{recovery}$ under various glucose conditions under two representative ODs.**

**(A-B)**, Linear fitting of $T_{recovery}$ data during M phase (blue lines) and D phase (red lines) under various glucose conditions with the initial OD of 0.5 (**A**) and 0.15 (**B**). Data condition, linear fitting slope, and $R^2$ are presented in the up left corner of each panel. Cell numbers for each data point are listed in Table S2.

**(C-D)** Comparison of $T_{recovery}$ for quiescent cells at day6 under different OD (**C**) and different glucose concentrations with two representative ODs (**D**). Cell numbers are indicated.

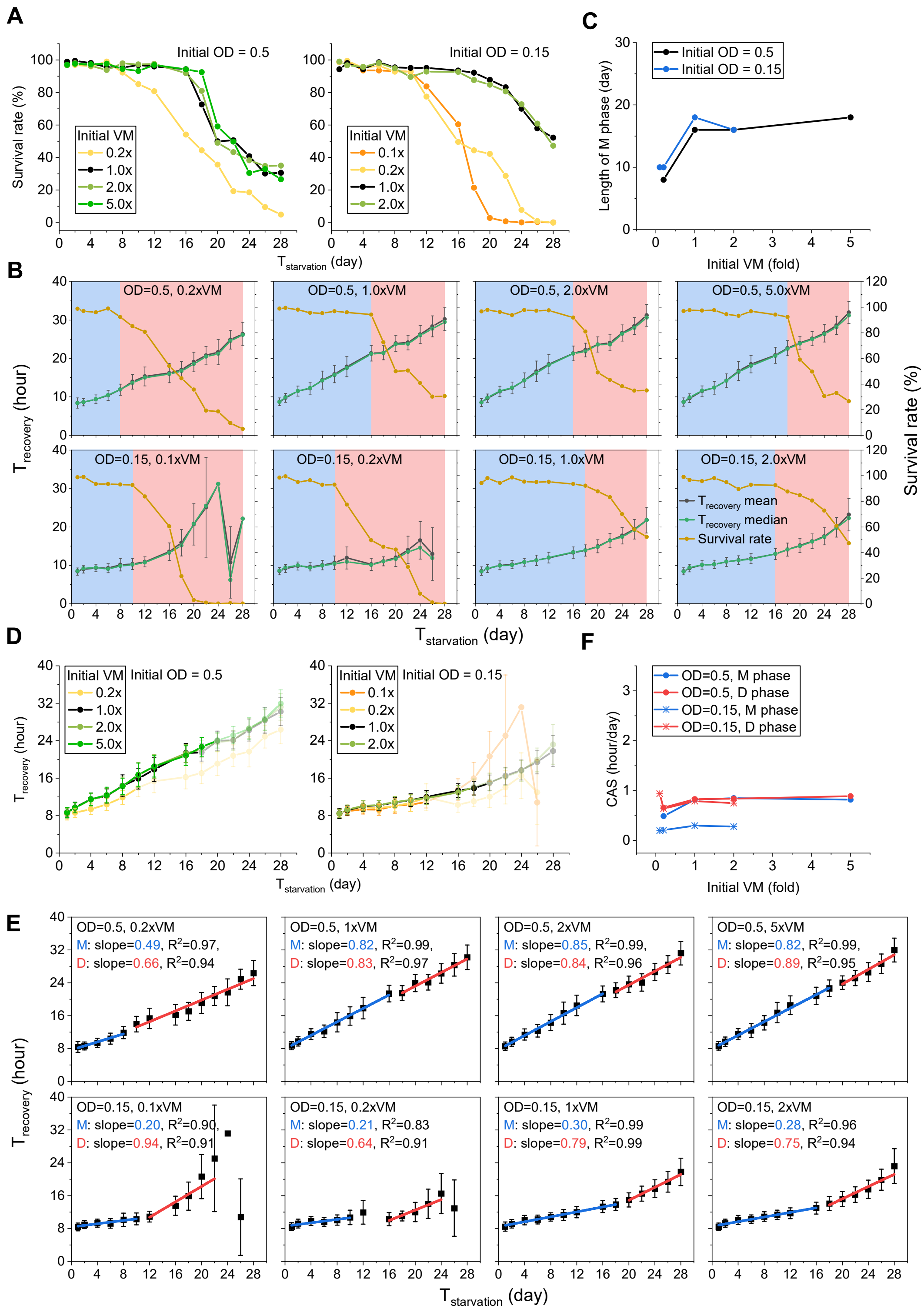


**Figure S4. Single cell aging behaviors under various VM conditions under two representative ODs.**
**(A-D)** Measured survival rate (**A**), M phase and D phase (**B**), length of M phase (**C**), and $T_{recovery}$ (**D**) under various vitamin and mineral concentrations (VM) under two representative ODs**.**
**(E)** Linear fitting of $T_{recovery}$ data in **D** for each condition during M phase (blue lines) and D phase (red lines). Data condition, linear fitting slope, and $R^2$ are presented in the up left corner of each panel.
(**F**) Comparison of CAS measured in **E**. Cell numbers for each data point are listed in Table S3.

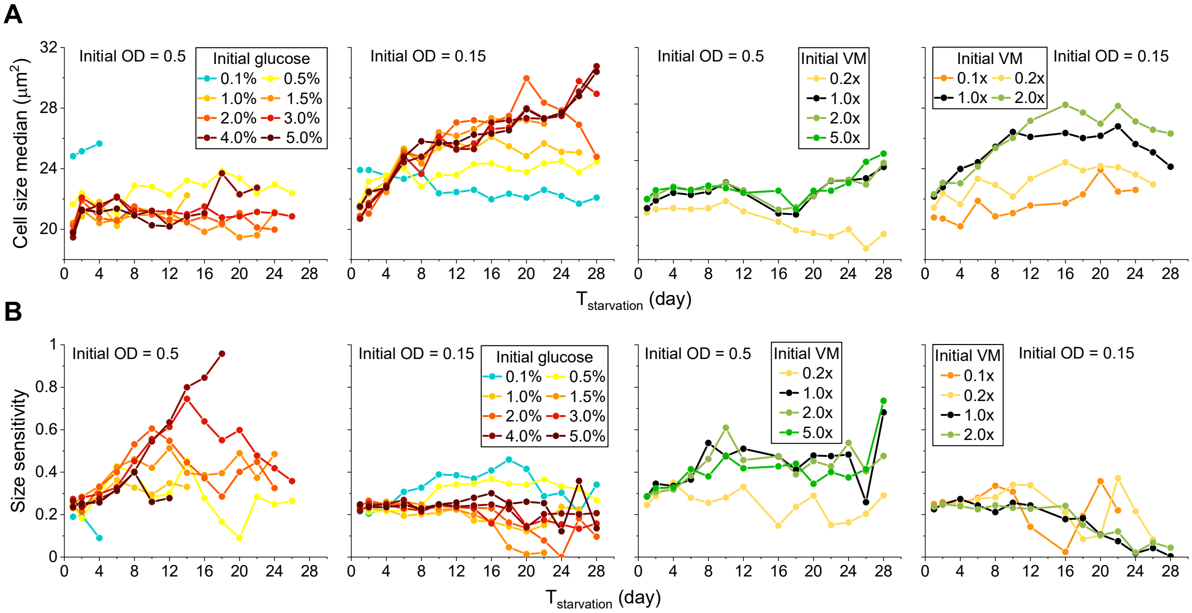


**Figure S5. Comparison of quiescent cell size and size sensitivity under various glucose and VM conditions.** **(A-B)** The change of quiescent cell size (**A**) and size sensitivity (**B**) of $Q_{-N}$ cells under various glucose and VM conditions under the initial OD of 0.5 and 0.15. Cell size in **A** is presented by the median value and data condition is presented in each panel. Cell numbers are listed in Table S2 and S3.

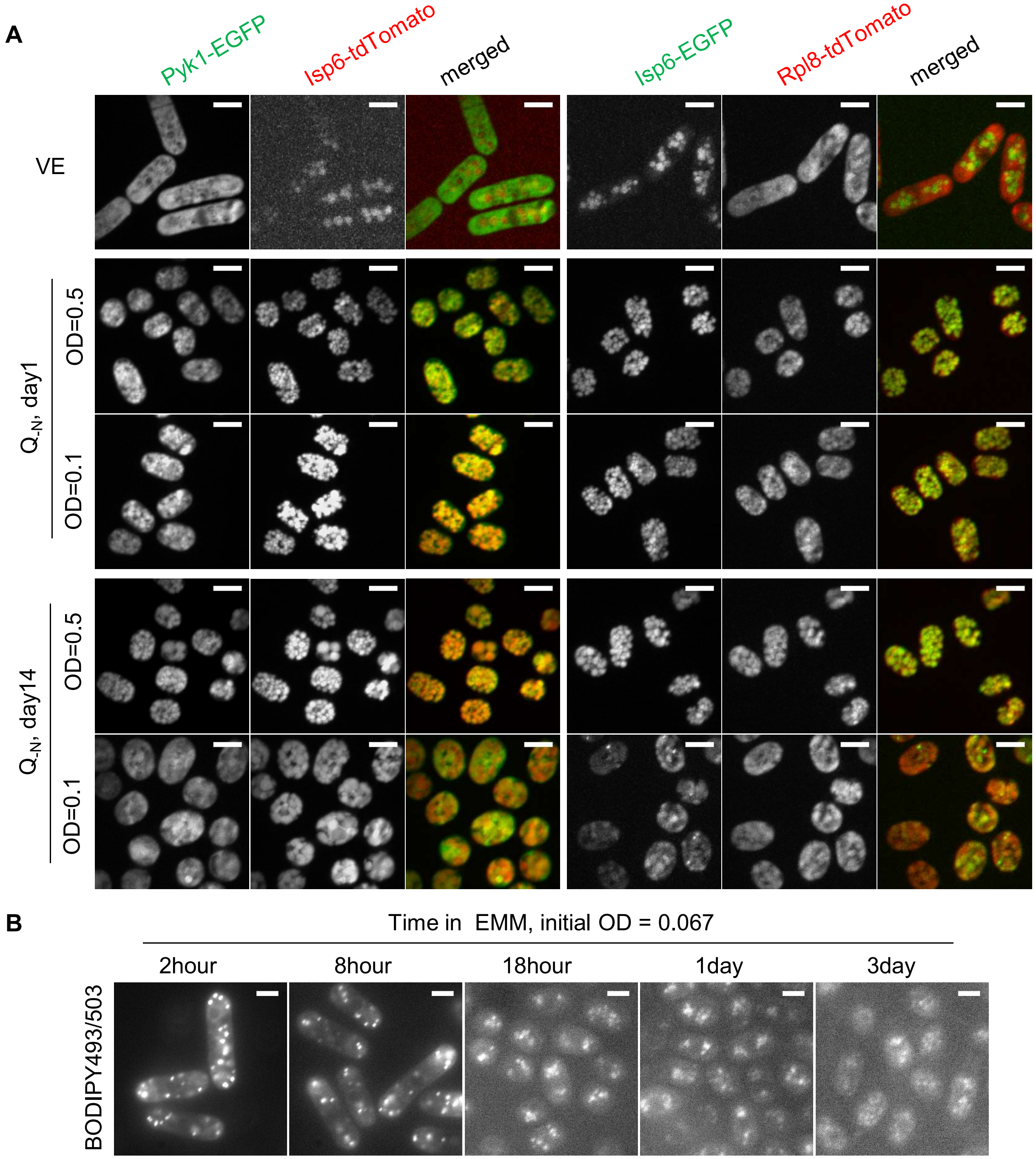


**Figure S6. The co-localization of glycolysis enzyme and ribosome with vacuolar, decrease of lipids accumulation at stationary phase.**

**(A)** The localization of Pyk1-EGFP, Isp6-tdTomato, Isp6-EGFP, and Rpl8-tdTomato in day1 and day14 $Q_{-N}$ cells under the initial OD of 0.5 and 0.1.

**(B)** Change of lipids accumulation during ST cell ageing. Lipids are stained with BODIPY493/503. The scale bar is 5μm.
