## supplemental tables S1-S7 for "Initial niche condition determines the aging speed and regenerative activity of quiescent cells"

**Supplementary Table S1. Analyzed numbers of single quiescent cells with various initial OD from imaging data**

$Q_{-N}$: nitrogen-starvation induced quiescence,
ST: stationary phase,
ND: no data

| Quiescence | Initial OD | Group | $T_{starvation}$ (day) | | | | | | | |
|---|---|---|---|---|---|---|---|---|---|---|
| | | | 1 | 2 | 4 | 6 | 8 | 10 | 12 | 14 |
| $Q_{-N}$ | 0.15 | total | 142 | 125 | 164 | 162 | 168 | ND | 184 | 181 |
| | | recovered | 141 | 123 | 157 | 162 | 164 | ND | 175 | 172 |
| | 0.3 | total | 127 | 121 | 156 | 179 | 180 | ND | 169 | 151 |
| | | recovered | 126 | 120 | 154 | 177 | 174 | ND | 163 | 149 |
| | 0.4 | total | 163 | 115 | 142 | 151 | 139 | 148 | 173 | 194 |
| | | recovered | 161 | 112 | 139 | 144 | 136 | 145 | 163 | 188 |
| | 0.5 | total | 156 | 172 | 127 | 184 | 152 | 152 | 190 | 161 |
| | | recovered | 152 | 171 | 124 | 180 | 148 | 147 | 185 | 154 |
| | 0.6 | total | 154 | 141 | 140 | 157 | 174 | 146 | 168 | 184 |
| | | recovered | 152 | 139 | 137 | 151 | 171 | 144 | 158 | 146 |
| | 0.8 | total | 162 | 156 | 138 | 167 | 174 | 156 | 192 | 292 |
| | | recovered | 160 | 155 | 137 | 161 | 166 | 138 | 133 | 174 |
| | 1.0 | total | 158 | 165 | 135 | 176 | 177 | 182 | 204 | 217 |
| | | recovered | 152 | 159 | 130 | 166 | 159 | 131 | 159 | 167 |
| ST | ST | total | 180 | 151 | 136 | 272 | 518 | 380 | 532 | ND |
| | | recovered | 180 | 132 | 96 | 96 | 62 | 12 | 3 | ND |

### Supplementary Table S2. Analyzed numbers of single $Q_{-N}$ cells with various initial glucose from imaging data

| Initial OD | Initial glucose | Group | $T_{starvation}$ (day) | | | | | | | | | | | | | | |
|---|---|---|---|---|---|---|---|---|---|---|---|---|---|---|---|---|---|
| | | | 1 | 2 | 4 | 6 | 8 | 10 | 12 | 14 | 16 | 18 | 20 | 22 | 24 | 26 | 28 |
| 0.5 | 0.1% | total | 127 | 164 | 164 | 148 | ND | ND | ND | ND | ND | ND | ND | ND | ND | ND | ND |
| | | recovered | 126 | 143 | 18 | 1 | ND | ND | ND | ND | ND | ND | ND | ND | ND | ND | ND |
| | 0.5% | total | 139 | 130 | 131 | 135 | 174 | 147 | 188 | 218 | 298 | 180 | 244 | 415 | 1444 | 1805 | ND |
| | | recovered | 138 | 127 | 127 | 133 | 103 | 45 | 54 | 58 | 56 | 23 | 18 | 29 | 47 | 21 | ND |
| | 1.0% | total | 115 | 153 | 126 | 168 | 155 | 136 | 175 | 310 | 277 | ND | ND | ND | ND | ND | ND |
| | | recovered | 114 | 150 | 125 | 158 | 131 | 117 | 161 | 66 | 0 | ND | ND | ND | ND | ND | ND |
| | 1.5% | total | 131 | 192 | 177 | 148 | 166 | 127 | 126 | 216 | 252 | 162 | 153 | 217 | 465 | ND | ND |
| | | recovered | 127 | 185 | 173 | 144 | 161 | 122 | 118 | 202 | 230 | 148 | 127 | 131 | 203 | ND | ND |
| | 2.0% | total | 147 | 158 | 174 | 127 | 163 | 162 | 139 | 178 | 155 | 184 | 165 | 336 | 404 | ND | ND |
| | | recovered | 144 | 156 | 168 | 125 | 158 | 155 | 128 | 167 | 141 | 158 | 141 | 201 | 243 | ND | ND |
| | 3.0% | total | 164 | 150 | 165 | 121 | 144 | 163 | 173 | 179 | 167 | 211 | 235 | 342 | 436 | 485 | ND |
| | | recovered | 162 | 142 | 161 | 119 | 143 | 158 | 159 | 161 | 138 | 168 | 177 | 216 | 238 | 209 | ND |
| | 4.0% | total | 177 | 164 | 178 | 140 | 153 | 154 | 156 | 201 | 212 | 248 | 305 | 770 | ND | ND | ND |
| | | recovered | 169 | 163 | 172 | 137 | 148 | 153 | 141 | 148 | 91 | 26 | 2 | 1 | ND | ND | ND |
| | 5.0% | total | 196 | 171 | 131 | 164 | 190 | 141 | 150 | 237 | ND | 246 | ND | ND | ND | ND | ND |
| | | recovered | 193 | 169 | 130 | 158 | 183 | 129 | 98 | 2 | ND | 0 | ND | ND | ND | ND | ND |
| 0.15 | 0.1% | total | 177 | 137 | 178 | 156 | 192 | 160 | 171 | 192 | 186 | 171 | 187 | 204 | 247 | 223 | 243 |
| | | recovered | 175 | 134 | 173 | 151 | 186 | 158 | 169 | 181 | 176 | 162 | 174 | 180 | 188 | 132 | 101 |
| | 0.5% | total | 210 | 143 | 191 | 192 | 177 | 186 | 179 | 188 | 185 | 172 | 188 | 169 | 178 | 174 | 225 |
| | | recovered | 206 | 140 | 185 | 185 | 169 | 178 | 172 | 182 | 177 | 169 | 183 | 161 | 169 | 166 | 212 |
| | 1.0% | total | 209 | 186 | 183 | 149 | 152 | 193 | 157 | 209 | 179 | 201 | 194 | 198 | 283 | 300 | ND |
| | | recovered | 208 | 180 | 182 | 142 | 146 | 182 | 147 | 205 | 175 | 184 | 174 | 167 | 214 | 201 | ND |
| | 1.5% | total | 183 | 144 | 163 | 183 | 145 | 142 | 164 | 164 | 172 | 244 | 206 | 284 | ND | ND | ND |
| | | recovered | 180 | 143 | 162 | 176 | 134 | 136 | 154 | 147 | 161 | 200 | 143 | 179 | ND | ND | ND |
| | 2.0% | total | 226 | 169 | 145 | 172 | 179 | 149 | 182 | 178 | 185 | 263 | 169 | 215 | 318 | 333 | 413 |
| | | recovered | 222 | 166 | 140 | 164 | 173 | 142 | 166 | 170 | 166 | 225 | 135 | 155 | 203 | 195 | 158 |
| | 3.0% | total | 206 | 157 | 150 | 169 | 188 | 161 | 179 | 183 | 221 | 184 | 205 | 179 | 269 | 302 | 404 |
| | | recovered | 203 | 154 | 147 | 161 | 182 | 151 | 174 | 176 | 209 | 172 | 173 | 155 | 211 | 192 | 183 |
| | 4.0% | total | 199 | 165 | 167 | 185 | 241 | 160 | 154 | 211 | 196 | 199 | 175 | 225 | 262 | 310 | 412 |
| | | recovered | 191 | 162 | 163 | 176 | 229 | 153 | 151 | 200 | 181 | 181 | 155 | 178 | 189 | 179 | 184 |
| | 5.0% | total | 185 | 132 | 141 | 167 | 253 | 164 | 199 | 255 | 219 | 256 | 188 | 237 | 219 | 284 | 398 |
| | | recovered | 185 | 131 | 139 | 164 | 241 | 158 | 190 | 235 | 203 | 230 | 170 | 180 | 146 | 160 | 160 |

ND: no data

### Supplementary Table S3. Analyzed numbers of single $Q_{-N}$ cells with various initial VM from imaging data

| Initial OD | Initial VM | Group | $T_{starvation}$ (day) | | | | | | | | | | | | | | |
|---|---|---|---|---|---|---|---|---|---|---|---|---|---|---|---|---|---|
| | | | **1** | **2** | **4** | **6** | **8** | **10** | **12** | **14** | **16** | **18** | **20** | **22** | **24** | **26** | **28** |
| 0.5 | 0.2x | total | 159 | 166 | 254 | 197 | 220 | 202 | 156 | ND | 266 | 639 | 464 | 475 | 376 | 470 | 426 |
| | | recovered | 157 | 161 | 244 | 195 | 203 | 171 | 126 | ND | 143 | 284 | 171 | 92 | 70 | 45 | 21 |
| | 1.0x | total | 185 | 177 | 210 | 182 | 149 | 158 | 176 | ND | 228 | 296 | 223 | 249 | 228 | 242 | 307 |
| | | recovered | 183 | 176 | 206 | 174 | 142 | 153 | 169 | ND | 215 | 215 | 111 | 126 | 93 | 73 | 92 |
| | 2.0x | total | 153 | 201 | 165 | 195 | 188 | 146 | 166 | ND | 259 | 300 | 277 | 277 | 261 | 250 | 329 |
| | | recovered | 148 | 196 | 159 | 183 | 184 | 142 | 162 | ND | 238 | 243 | 136 | 120 | 100 | 87 | 115 |
| | 5.0% | total | 203 | 213 | 189 | 171 | 164 | 252 | 162 | ND | 190 | 306 | 274 | 246 | 263 | 237 | 259 |
| | | recovered | 197 | 208 | 184 | 168 | 155 | 232 | 157 | ND | 180 | 283 | 162 | 123 | 80 | 78 | 69 |
| 0.15 | 0.1x | total | 254 | 216 | 234 | 185 | 263 | 224 | 257 | ND | 501 | 695 | 457 | 523 | 797 | 780 | 1040 |
| | | recovered | 251 | 213 | 219 | 173 | 244 | 205 | 216 | ND | 301 | 149 | 13 | 4 | 1 | 3 | 1 |
| | 0.2x | total | 203 | 192 | 245 | 141 | 181 | 257 | 182 | ND | 256 | 360 | 322 | 425 | 410 | 979 | 662 |
| | | recovered | 200 | 191 | 233 | 137 | 168 | 238 | 141 | ND | 127 | 160 | 136 | 122 | 32 | 9 | 0 |
| | 1.0x | total | 228 | 209 | 214 | 141 | 174 | 164 | 186 | ND | 169 | 179 | 173 | 215 | 216 | 341 | 358 |
| | | recovered | 214 | 204 | 202 | 139 | 166 | 156 | 177 | ND | 158 | 165 | 151 | 178 | 152 | 197 | 187 |
| | 2.0x | total | 199 | 184 | 210 | 160 | 156 | 172 | 224 | ND | 163 | 219 | 218 | 218 | 253 | 276 | 238 |
| | | recovered | 197 | 178 | 201 | 157 | 148 | 154 | 208 | ND | 151 | 192 | 184 | 175 | 184 | 168 | 113 |

ND: no data

**Supplementary Table S4. Fission yeast strains used in this study**

| Strain | Genotype | Reference |
|---|---|---|
| 972h⁻ | *972 h⁻* | Gift from Dr. Dao-chun Kong |
| LQF001 | *972 h⁻ Cdc13-yEGFP-$T_{yADH1}$-kanMX6 Rum1-tdTomato-$T_{Rum1}$-natMX6* | This study |
| LQF006 | *972 h⁻ Aco1-yEGFP-$T_{yADH1}$-kanMX6 Hta2-tdTomato-$T_{yADH1}$-natMX6* | This study |
| LQF068 | *972 h⁻ Pyk1-yEGFP-$T_{yADH1}$-kanMX6 Rpl8-tdTomato-$T_{yADH1}$-natMX6* | This study |
| LQF069 | *972 h⁻ Isp6-yEGFP-$T_{yADH1}$-kanMX6 Dga1-tdTomato-$T_{yADH1}$-natMX6* | This study |
| LQF072 | *972 h⁻ Pyk1-yEGFP-$T_{yADH1}$-kanMX6 Isp6-tdTomato-$T_{yADH1}$-natMX6* | This study |
| LQF079 | *972 h⁻ Isp6-yEGFP-$T_{yADH1}$-kanMX6 Rpl8-tdTomato-$T_{yADH1}$-natMX6* | This study |

### Supplementary Table S5. Plasmids used in this study

| Plasmid name | Genotype | Reference |
|---|---|---|
| pDH3 | *pCFP-T*$_{yADH1}$*-kanMX6* | Gift from Dr. Li-lin Du |
| pMD19 | *pMD19* | TaKaRa Cat# 3271 |
| pFA6a-natMX6 | *pFA6a-natMX6* | Lab stock |
| PP001 | *pCFP-natMX6* | This study |
| PP002 | *pNI8T-whi5-yEGFP* | Lab stock |
| PP003 | *pNI8T-tdTomato* | Lab stock |
| PP004 | *pyEGFP-T*$_{yADH}$*1-kanMX6* | This study |
| PP005 | *ptdTomato-T*$_{yADH1}$*-natMX6* | This study |
| PP006 | *pMD19-Leftleg*$_{Rum1}$*-tdTomato-T*$_{yADH1}$*-natMX6-Rightleg*$_{Rum1}$ | This study |
| PP007 | *pMD19-T*$_{Rum1}$ | This study |
| PP008 | *pMD19-Leftleg*$_{Rum1}$*-tdTomato-T*$_{Rum1}$*-natMX6-Rightleg*$_{Rum1}$ | This study |

**Supplementary Table S6. Oligonucleotides used in this study**

| Oligonucleotides | Sequence (5' – 3') |
|---|---|
| Cdc13-GK fusion, leftleg forward | TTTGGCCGTAAATATTATTG |
| Cdc13-GK fusion, leftleg reverse | AACCCGGGGATCCGTCGACCCCATTCTTCATCTTTCATGT |
| Cdc13-GK fusion, rightleg forward | CGAGCTCGAATTCATCGATTTAGTGTATTGTGCATATCACT |
| Cdc13-GK fusion, rightleg reverse | TTCACCTCAAGGAACTATCA |
| Rum1 3'UTR T-cloning forward | GGCGCGCCCTTTTTTTCGCATTTGTAATTGT |
| Rum1 3'UTR T-cloning reverse | AGATCTCTAGAATAGTATTGTATATGCGGT |
| Rum1-TN fusion, leftleg forward | GATAGATGAAATCCCTGAAAG |
| Rum1-TN fusion, leftleg reverse | ACCCGGGGATCCGTCGACCTCGTAATAAATTGTGCCTGTT |
| Rum1-TN fusion, leftleg forward | CGAGCTCGAATTCATCGATCTTTTTTTCGCATTTTGTAATTG |
| Rum1-TN fusion, leftleg reverse | CTATAGTTTCCAAACCAGCAA |
| Aco1- GK fusion, leftleg forward | CTCGTTATTTGGGAGGCG |
| Aco1- GK fusion leftleg reverse | CCCGGGGATCCGTCGACCCTTTTGCTTGTGCATGTTTG |
| Aco1-GK fusion, rightleg forward | CGAGCTCGAATTCATCGATAGGACTTATAAGCCTTCCGG |
| Aco1- GK fusion, rightleg reverse | GCAAGACTGTAGTTTGCTC |
| Hta2-TN fusion, leftleg forward | GTCGGTCGTGTTCATCGTTT |
| Hta2-TN fusion, leftleg reverse | AACCCGGGGATCCGTCGACCAAGCTCTTGGCTAGGCTTGC |
| Hta2-TN fusion, rightleg forward | ACGAGCTCGAATTCATCGATACTCTTAAATGAATGATAAATTGT |
| Hta2-TN fusion, rightleg reverse | AAAGGGAAGAGGAAAACACG |
| Pyk1-GK fusion, leftleg forward | TTGAAGCCGTTACCTACATG |
| Pyk1-GK fusion, leftleg reverse | CCCGGGGATCCGTCGACCCTCGGCAACGGTAAGACG |
| Pyk1-GK fusion, rightleg forward | CGAGCTCGAATTCATCGATGCTGATTATGACGATCCAGT |
| Pyk1-GK fusion, rightleg reverse | TTTTCCAATTGCTGAGATAC |
| Rpl8-TN fusion, leftleg forward | AAAGAAGCAACGTTTGGTTG |
| Rpl8-TN fusion, leftleg reverse | CCCGGGGATCCGTCGACCGAGACGAACAGTAGCGGCAG |
| Rpl8-TN fusion, rightleg forward | CGAGCTCGAATTCATCGATATAAATTCATCTGGATTGAAAATG |
| Rpl8-TN fusion, rightleg reverse | TATTCGTTTGGGTCTCCTTG |
| Isp6-TN/GK fusion, leftleg forward | GCTGCTGGAAATGAGTATGA |
| Isp6-TN/GK fusion, leftleg reverse | CCCGGGGATCCGTCGACCTTCTTGAGCACCATTGAAAG |
| Isp6-TN/GK fusion, rightleg forward | CGAGCTCGAATTCATCGATATTTTACTTCATGTTGGATGCATT |
| Isp6-TN/GK fusion, rightleg reverse | TTGCATAACAGCGGCACA |

GK: *yEGFP-*$T_{yADH1}$*-kanMX6* cassette,
TN: *tdTomato-*$T_{yADH1}$*-natMX6* cassette,
leftleg: homology DNA targeting left half of recombination site,
rightleg: homology DNA targeting right half of the recombination site.

### Supplementary Table S7. Synthetic media used in this study

| Category | Variants | Final concentration of adjusted component (per liter) | | | |
|---|---|---|---|---|---|
| | | $NH_4Cl$ | glucose | 1000x Vitamin stock | 10000x Mineral stock |
| EMM | Standard | 5.0g | 20g | 1mL | 0.1mL |
| EMM-C | Standard | 5.0g | 0g | 1mL | 0.1mL |
| EMM-N | Standard | 0g | 20g | 1mL | 0.1mL |
| | 0.1% glucose | 0g | 1g | 1mL | 0.1mL |
| | 0.5% glucose | 0g | 5g | 1mL | 0.1mL |
| | 1.0% glucose | 0g | 10g | 1mL | 0.1mL |
| | 1.5% glucose | 0g | 15g | 1mL | 0.1mL |
| | 2.0% glucose | 0g | 20g | 1mL | 0.1mL |
| | 3.0% glucose | 0g | 30g | 1mL | 0.1mL |
| | 4.0% glucose | 0g | 40g | 1mL | 0.1mL |
| | 5.0% glucose | 0g | 50g | 1mL | 0.1mL |
| | 0.1x VM | 0g | 20g | 0.1mL | 0.01mL |
| | 0.2x VM | 0g | 20g | 0.2mL | 0.02mL |
| | 1.0x VM | 0g | 20g | 1mL | 0.1mL |
| | 2.0x VM | 0g | 20g | 2mL | 0.2mL |
| | 5.0x VM | 0g | 20g | 5mL | 0.5mL |
| EMM-N-C | EMM-N-C | 0g | 0g | 1mL | 0.1mL |

**Captions for Supplementary Movies S1 to S5**

**Movie S1.**
Nitrogen-starved quiescent cells accumulate LD during quiescence establishment.

**Movie S2.**
Proliferating cells store basal level LD.

**Movie S3.**
Recovery of $Q_{-N}$ cells starved for 16 days under initial OD=0.5 with 0.5% initial glucose.

**Movie S4.**
Recovery of Q-N cells starved for 16 days under initial OD=0.5 with 2.0% initial glucose.

**Movie S5.**
Recovery of Q-N cells starved for 16 days under initial OD=0.5 with 4.0% initial glucose.
